# Optimal control of combination immunotherapy for a virtual murine cohort in a glioblastoma-immune dynamics model

**DOI:** 10.1101/2024.04.29.591725

**Authors:** Hannah G. Anderson, Gregory P. Takacs, Jeffrey K. Harrison, Libin Rong, Tracy L. Stepien

## Abstract

The immune checkpoint inhibitor anti-PD-1, commonly used in cancer immunotherapy, has not been successful as a monotherapy for the highly aggressive brain cancer glioblastoma. However, when used in conjunction with a CC-chemokine receptor-2 (CCR2) antagonist, anti-PD-1 has shown efficacy in preclinical studies. In this paper, we aim to optimize treatment regimens for this combination immunotherapy using optimal control theory. We extend a treatment-free glioblastoma-immune dynamics ODE model to include interventions with anti-PD-1 and the CCR2 antagonist. An optimized regimen increases the survival of an average mouse from 32 days post-tumor implantation without treatment to 111 days with treatment. We scale this approach to a virtual murine cohort to evaluate mortality and quality of life concerns during treatment, and predict survival, tumor recurrence, or death after treatment. A parameter identifiability analysis identifies five parameters suitable for personalizing treatment within the virtual cohort. Sampling from these five practically identifiable parameters for the virtual murine cohort reveals that personalized, optimized regimens enhance survival: 84% of the virtual mice survive to day 100, compared to 60% survival in a previously studied experimental regimen. Subjects with high tumor growth rates and low T cell kill rates are identified as more likely to die during and after treatment due to their compromised immune systems and more aggressive tumors. Notably, the MDSC death rate emerges as a long-term predictor of either disease-free survival or death.

**Highlights:**

- A mathematical model of glioma-immune dynamics integrates combination immunotherapy.
- An optimized regimen extends survival in an average virtual mouse by 79 days.
- Quality of life and survival outcomes were evaluated for a virtual murine cohort.
- A high death rate of myeloid-derived suppressor cells predicts long-term survival.

## 1. Introduction

Approximately 350,000 people are newly diagnosed with brain tumors across the globe each year, with about 250,000 deaths worldwide (Ilic and Ilic (2023)[Fig. 1]). Of these brain tumors, glioblastoma (GBM) is the most aggressive and most common type–comprising 49% of all primary brain malignancies (Schaff and Mellinghoff (2023)). People diagnosed with GBM start to experience symptoms such as worsening headaches, seizures, memory loss or confusion, and unsteadiness (Grant (2004); Gilard et al. (2021)). The current standard of care is surgical resection followed by radiotherapy and chemotherapy with temozolomide (TMZ), which has a median survival of 14.6 months (Stupp et al. (2005)).

**Figure 1:**
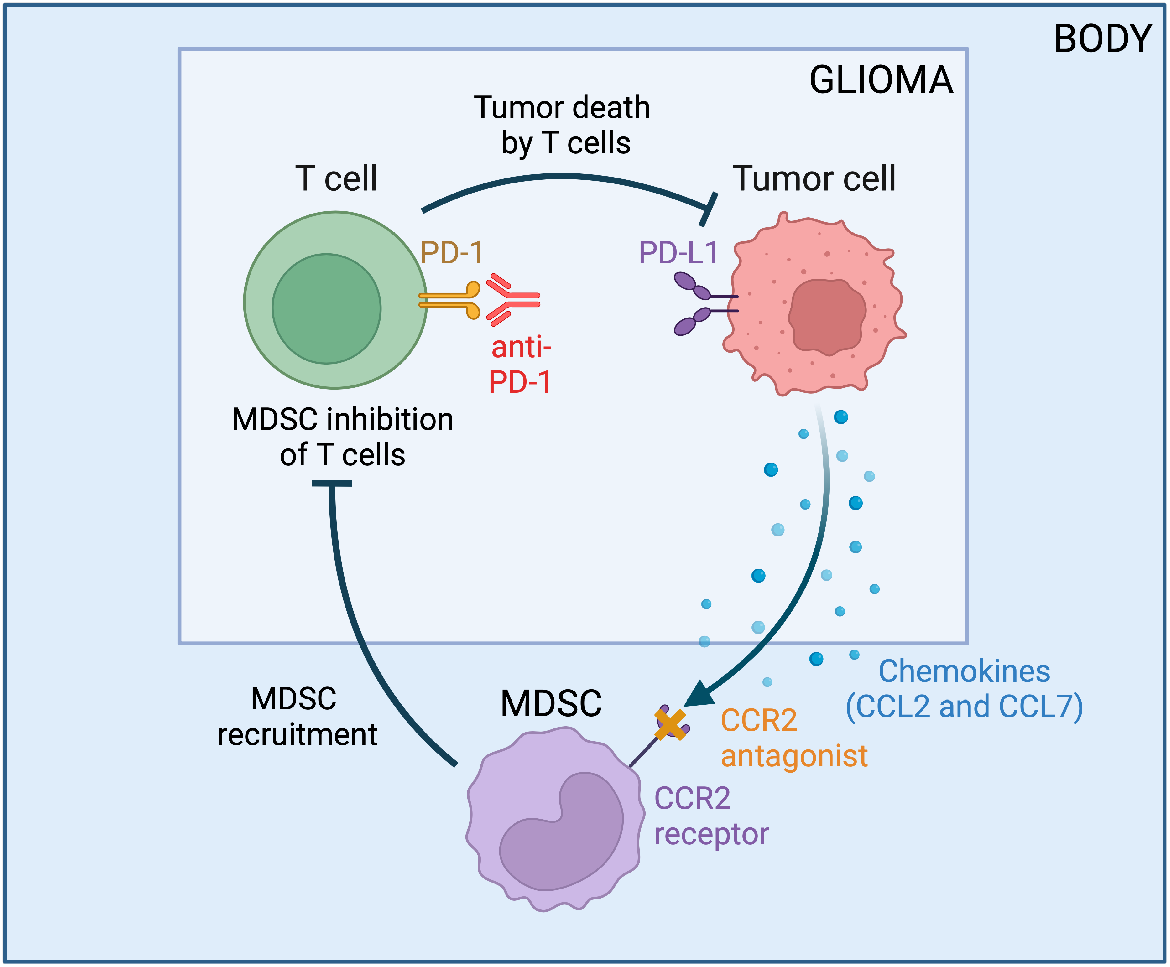
Tumor-immune interactions in glioblastoma when treated with anti-PD-1 and a CCR2 antagonist. Flowchart created with BioRender.com.

Preclinical studies and clinical trials are testing new therapies for GBM with the aim to increase this survival time. One such therapy is a combination of two immunotherapies: the immune checkpoint inhibitor, anti-PD-1, and a CC-chemokine receptor-2 (CCR2) antagonist that prevents the tumorinduced recruitment of immunosuppressive myeloid-derived suppressor cells (MDSCs). These treatments have been tested as monotherapies and combination therapies in other cancers, such as renal cell carcinoma and pancreatic ductal adenocarcinoma, and hold promise for GBM (Orth et al. (2019); Choueiri et al. (2022); Flores-Toro et al. (2020)).

Anti-PD-1 is a monoclonal antibody designed to inhibit the PD-1 receptor of T cells in order to elicit an immune response to cancer. PD-1 inhibitors, such as nivolumab and pembrolizumab, are currently approved for the treatment of melanoma, non-small cell lung cancer (NSCLC), renal cell carcinoma, squamous cell carcinoma of the head and neck, metastatic colorectal cancer, as well as urothelial, esophageal, and gastric cancers, among others (Lee et al. (2022)).

Anti-PD-1 causes fewer adverse events than many other immune checkpoint inhibitors (Tanaka and Okamura (2013)); however, it is not without its safety concerns. Common side effects include fatigue, rash, nausea, weakness, shortness of breath, constipation, vomiting, headache, and fever (Martins et al. (2019)). About 10% of patients treated with anti-PD-1 experience grade 3 or higher immune-related adverse events (irAEs) (Martins et al. (2019)), where irAEs are graded on a scale of 1 to 5 (1 being mild, and 5 fatal) (Kumar et al. (2017); Abdel-Wahab et al. (2017)). Anti-PD-1 is often used concurrently with another treatment, so given its toxicity profile, it is shrewd to combine it with a therapeutic exhibiting fewer irAEs, such as a CCR2 antagonist.

A CCR2 antagonist is a small molecule which targets the CCL2/CCR2 signaling pathway to prevent the recruitment of CCR2-expressing monocytes and macrophages to the tumor site (Fei et al. (2021)). CCR2-expressing cells like myeloid-derived suppressor cells (MDSCs) aid in immune suppression at the tumor site (Takacs et al. (2021)). CCR2 antagonists (such as BMS813160, PF-04136309, and CCX872) are in clinical trials for pancreatic ductal adenocarcinoma (ClinicalTrials.gov [Internet]. Sidney Kimmel Comprehensive Cancer Center at Johns Hopkins (2019 Dec 12 -)) and advanced renal cell carcinoma (ClinicalTrials.gov [Internet]. Bristol-Myers Squibb (2017 Feb 2 - 2021 Nov 23)), where both trials use anti-PD-1 concurrently. CCR2^+^ MDSCs are found within the tumor microenvironment of glioblastomas (Takacs et al. (2021)), making CCR2 inhibition a desirable therapeutic mechanism for exploration.

CCR2 antagonists are relatively safe, so this immunotherapy is a suitable option to combine with anti-PD-1. In a clinical trial for pancreatic ductal adenocarcinoma, combination treatment with the CCR2 antagonist BMS813160 and anti-PD-1 was determined to be safe with only one grade 3 or higher adverse event in a single patient (Christenson et al. (2023)).

While both CCR2 inhibition and anti-PD-1 have failed as monotherapies for glioblastoma, preclinical murine models have shown that a CCR2 antagonist improves the efficacy of anti-PD-1 in cancers such as glioblastoma, bladder, and breast cancer (Flores-Toro et al. (2020); Tu et al. (2020)). Since the combination therapy is efficacious in preclinical models and tolerable in early phase clinical trials, we focus on optimizing this treatment regimen using optimal control theory.

The goal of optimal control is to find the “controls” for a dynamical system which allow us to reach a desired outcome by minimizing (or maximizing) an objective functional. Optimal control theory has been used to optimize policies and practices for a wide variety of industries including economics (Sheng et al. (2014); Wu et al. (2015); Li et al. (2019); Kaszkurewicz and Bhaya (2022)), aerospace technologies (Benson (2005); Longuski et al. (2014); Villegas Díaz et al. (2019); García-Heras et al. (2019); Sun et al. (2019); Zhao et al. (2019)), and the medical field (Martin and Teo (1994); Jung et al. (2002); Ledzewicz et al. (2013); Wang and Schättler (2016); Ledzewicz and Moore (2017); Ratajczyk et al. (2018); Moore et al. (2018); Fernández and Pola (2019); Ledzewicz et al. (2019); Gutiérrez-Diez and Russo (2020); Jarrett et al. (2020); Sharp et al. (2020); Rautela et al. (2023); Luo et al. (2023); Valega-Mackenzie et al. (2024)). Control theory was even used in the development of the “artificial pancreas,” which became FDA approved in 2016 for patients with type 1 diabetes (FDA (2016)). This method has been used to optimize oncology treatment (Luo et al. (2023)), reduce the spread of infectious diseases (Jung et al. (2002)), and address chronic health issues (Christodoulides et al. (2017)). Within oncology, researchers have used optimal control theory to improve the administration of various therapies, such as chemotherapy (Martin and Teo (1994)), immunotherapy (Ledzewicz et al. (2013)), and radiation (Jarrett et al. (2020)), and have addressed different situations in oncology including drug resistance (Rautela et al. (2023)), tumor heterogeneity (Wang and Schättler (2016)), combination therapy (Ratajczyk et al. (2018)), and treatment personalization (Gutiérrez-Diez and Russo (2020)). Optimal control theory exhibits incredible potential to improve cancer therapeutics by elucidating regimens which minimize both the tumor size and therapeutic dosages. These optimized regimens can decrease mortality rates while improving patients’ quality of life through the minimization of immune-related adverse events.

Many mathematical models of tumor-immune dynamics have been developed (Sardar et al. (2024); Cherraf et al. (2023); Song et al. (2021); Liu et al. (2021); Yin et al. (2019); Shariatpanahi et al. (2018); Nikolopoulou et al. (2018); Lai and Friedman (2017); Eladdadi et al. (2014); de Pillis et al. (2005); de Pillis and Radunskaya (2003)), including a few that focus specifically on GBM (Anderson et al. (2023); Santurio and Barros (2022); Khajanchi (2021); Storey et al. (2020)). The basis of our model stems from Lai and Friedman (2017), who developed a PDE model of general tumor-immune dynamics incorporating the PD-L1-PD-1 immune checkpoint and combination treatment with anti-PD-1 and a cancer vaccine. Nikolopoulou et al. (2018) then simplified it into an ODE model of a general tumor and T cells with anti-PD-1 treatment. Shariatpanahi et al. (2018) also developed an ODE model for the general tumor, but instead of incorporating the PD-L1-PD-1 immune checkpoint, they incorporated immune suppression via MDSCs and implemented treatment with a chemotherapy. By drawing from both Nikolopoulou et al. (2018) and Shariatpanahi et al. (2018), Anderson et al. (2023) developed a treatment-free ODE model including immune suppression via the PD-L1-PD1 complex and MDSCs and made it GBM-specific by estimating parameter distributions using data from glioma-bearing mice. Here, we extend the ODE model of Anderson et al. (2023), which incorporates tumor cells, T cells, and MDSCs, to include treatment with anti-PD-1 and a CCR2 antagonist and subsequently apply optimal control theory.

We optimize treatment for an average murine subject and then consider characteristics of the resulting regimen. The greater aim, however, is to scale this optimization to a virtual murine cohort of tens of thousands of subjects to determine personalized treatment regimens. During treatment, mice are categorized based on mortality or quality of life concerns due to tumor burden or elevated drug toxicities. After treatment, survival outcomes are determined by categorizing subjects in terms of disease-free and/or progression-free survival, tumor recurrence, and death. We obtain an estimate of the median survival and determine a subpopulation of subjects best suited for concurrent therapy with anti-PD-1 and a CCR2 antagonist.

The paper is organized as follows: in Section 2, we describe the GBMimmune dynamics model with the inclusion of the two treatments/controls (anti-PD-1 and the CCR2 antagonist), and then proceed to describe the objective functional to be minimized using optimal control theory in Section 3. In Section 4, we prepare for treatment personalization by performing parameter identifiability analysis of the treatment-free model. Treatment personalization results are presented in Section 6 followed by a discussion of the results in Section 8.

## 2. Model

We adapt the GBM-specific tumor-immune dynamics model from Anderson et al. (2023) to incorporate treatment with anti-PD-1 and a CCR2 antagonist. We consider the influence of these immunotherapies on the number of tumor cells, *C*, activated T cells, *T*, and myeloid-derived suppressor cells (MDSCs), *M*. Figure 1 displays a visual representation of the tumorimmune dynamics with treatment, and Table 1 lists a description of each parameter along with its units and summary statistics, and the resulting system of equations is

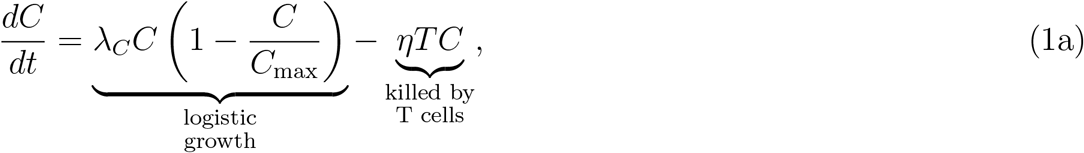

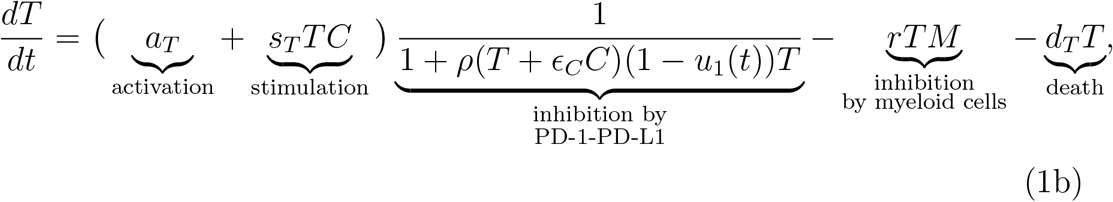

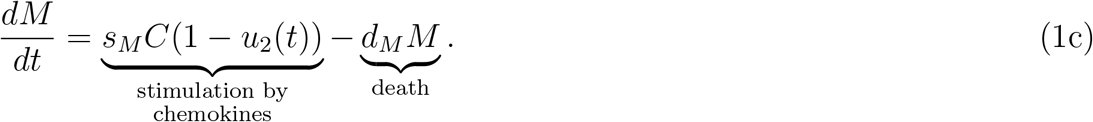

Cancer growth is represented logistically, where *λ*_*C*_ is the tumor growth rate, and *C*_max_ is the tumor carrying capacity. Upon interaction with T cells, cancer cells are killed at a rate of *η*.

**Table 1:**
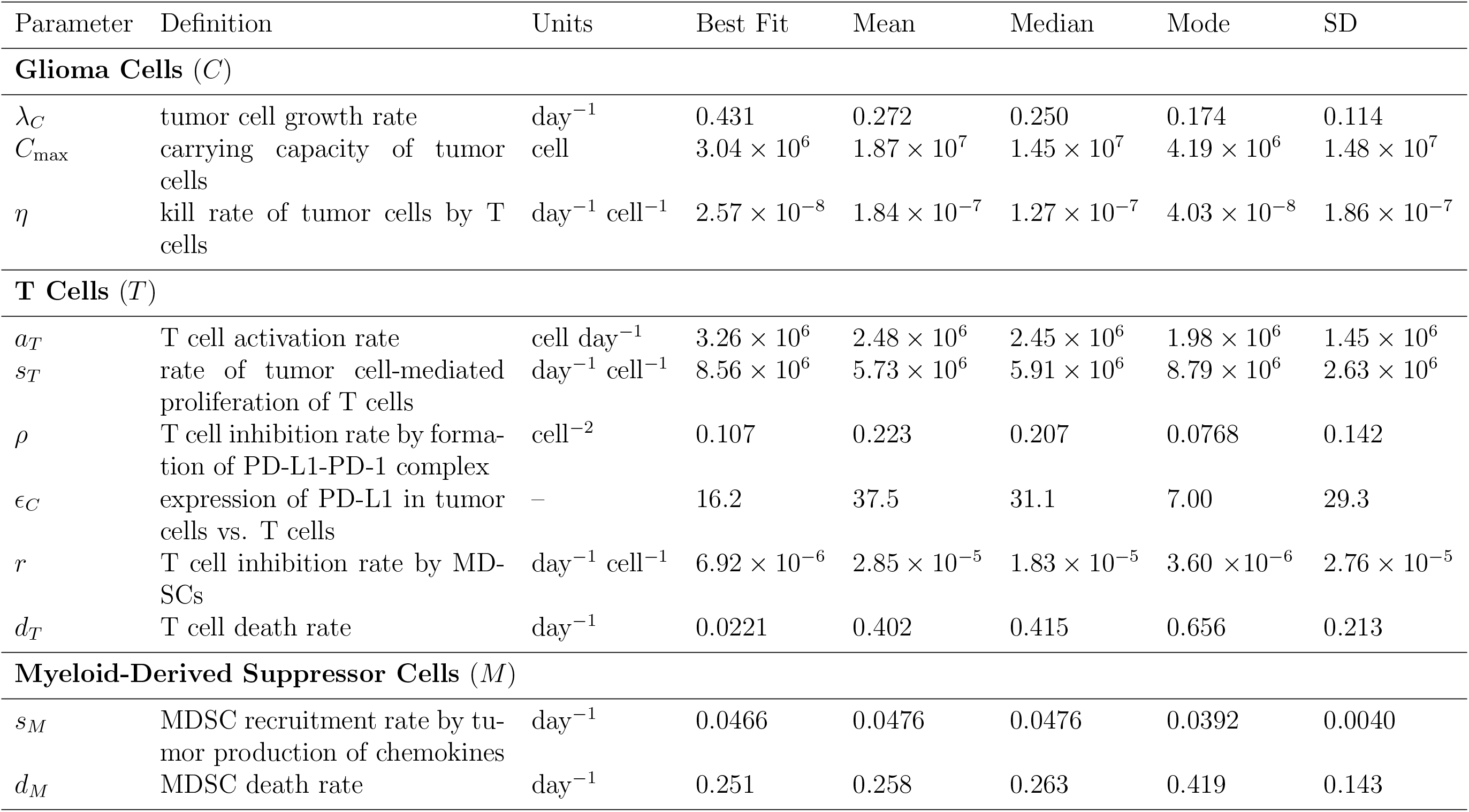
Model parameters of the glioblastoma (GBM)–immune dynamics model (1) without treatment were determined using the approximate Bayesian computation (ABC) rejection method in Anderson et al. (2023). The summary statistics of *s*_*M*_ were re-estimated here using the updated equation for MDSCs (1c).

T cell activation occurs at a rate of *a*_*T*_. The presence of a tumor stimulates an immune response, thus resulting in an influx of T cells to the tumor site at a rate of *s*_*T*_. Once at the tumor site, T cells are inhibited by the formation of PD-L1-PD-1. We assume that the interaction of PD-1 on T cells is proportional as *ρ*_1_*T*, and the interaction of PD-L1 on T cells and tumor cells is proportional as *ρ*_*L*1_(*T* +*ϵ*_*C*_*C*), where the level of tumor upregulation of PDL1 is represented by the parameter *ϵ*_*C*_. Anti-PD-1 (*u*_1_) binds to available PD1 on T cells, thus decreasing T cell inhibition by the PD-L1-PD-1 complex. Simplifying due to parameter non-identifiability, total formation of the PDL1-PD-1 complex is represented by

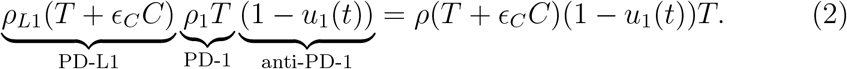

T cells are further inhibited by MDSCs at a rate of *r*, and they die naturally at a rate of *d*_*T*_.

MDSC recruitment to the tumor site is in part a response to glioma expression of chemokines CCL2 and CCL7, which are ligands of the CCR2 receptor expressed by MDSCs (Takacs et al. (2022)). A CCR2 antagonist (*u*_2_) decreases this recruitment by binding to the CCR2 receptor. Details of the derivation for the CCR2 antagonist term are given in Appendix A. We assume that MDSCs die naturally at a rate of *d*_*M*_. For simplicity, although we included splenic expansion of MDSCs in the model of Anderson et al. (2023), global sensitivity analysis using the eFAST method showed that the model was insensitive to that term, and thus, here in equation (1c), we do not include it. Aside from the inclusion of *u*_1_(*t*) and *u*_2_(*t*) in (1), this deletion was our only modification to the original Anderson et al. (2023) model.

We define *u*_1_(*t*) to be the percent reduction of the PD-L1-PD-1 inhibition rate (*ρ*) by anti-PD-1 at time *t* and *u*_2_(*t*) to be the percent reduction of the MDSC recruitment rate (*s*_*M*_) by the CCR2 antagonist. Both *u*_1_(*t*) and *u*_2_(*t*) are considered to be Lebesgue integrable functions for practical and numerical reasons, since this ensures that the percent reduction is nonnegative and that the total cumulative percent reduction is finite.

## 3. Formulation of Treatment as an Optimal Control Problem

We seek to minimize tumor burden, *C*, as well as toxicity of each immunotherapy, *u*_1_ and *u*_2_, to avoid immune-related adverse events (irAEs).

The optimal control problem can be stated as

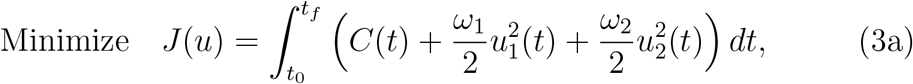

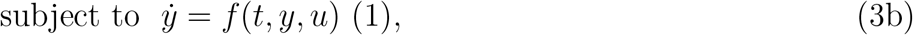

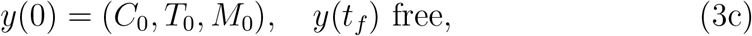

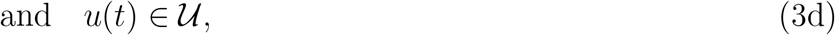

where *y* = (*C, T, M*) and *t*_0_ and *t*_*f*_ are the initial and final treatment time points, respectively. The space 𝒰 is defined as

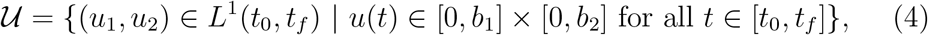

where *b*_1_ and *b*_2_ are the maximum percentage that anti-PD-1, *u*_1_, and the CCR2 antagonist, *u*_2_, can minimize PD-L1-PD-1 inhibition, *ρ*, and MDSC recruitment, *s*_*M*_, respectively, on the treatment interval [*t*_0_, *t*_*f*_ ].

In the objective functional (3a), the first term represents the cumulative amount of tumor cells. Each immunotherapy is modeled quadratically as convexity is mathematically advantageous for determining the minimum (Schättler and Ledzewicz (2015)). Weights *ω*_1_ and *ω*_2_ are functions of the tumor carrying capacity, *C*_max_, to balance the large tumor cell count with the considerably smaller dosage levels. Unlike the fairly tolerable toxicity profile of CCR2 antagonists (Tu et al. (2020)), immune checkpoint inhibitors such as anti-PD-1 are often associated with irAEs (Sandigursky and Mor (2018)). Therefore, we assume that *ω*_1_ *> ω*_2_. In Appendix B, we obtain the necessary conditions for optimality using Pontryagin’s minimum principle (Chiang (1999)[Ch. 7]), which addresses the Hamiltonian, adjoint equations, and transversality conditions to ensure there is a unique characterization of the optimal pair, (*u*_1_, *u*_2_).

## 4. Parameter Identifiability

In preparation for treatment personalization (Section 6), we perform parameter identifiability analysis (IA) to determine suitable parameters of interest (Guillaume et al. (2019); Miao et al. (2011)). By determining parameters which can be uniquely identified with data, this ensures that markers found for treatment failure or success in Section 6 can be used in practice. Errors in the model structure and the data are both sources of parameter non-identifiability, and these errors result in two categories of parameter identifiability: structural and practical. Structural (or *a priori*) IA offers a best-case scenario approach where the data set is assumed to be complete and noise-free and, thus, only the structure of the model affects the unique structural identification of parameters. Practical (or *a posteriori*) IA, on the other hand, offers a real-world approach where errors in data collection such as data noise and sparsity are taken into account.

We perform structural IA analytically using the differential algebra approach developed by Audoly et al. (2001). An overview of the method and its application to our model is stated in Appendix C. Results show that the treatment-free model is globally structurally identifiable with respect to tumor cell count data. This means that all parameters can be uniquely identified given a completely thorough and accurate data set on the tumor cell population. Thus, the model structure does not need to be modified.

We numerically calculate practical IA in Appendix D using the Fisher information matrix and profile likelihoods with murine data from Anderson et al. (2023). Analysis of the Fisher information matrix indicates that at most 5 parameters are practically identifiable, and sensitivity analysis identified the 6 most sensitive parameters. Profile likelihoods of the sensitive parameters showed that 5 are practically identifiable: tumor growth rate (*λ*_*C*_), T cell kill rate (*η*), inhibition rates by PD-L1-PD-1 (*ρ*) and by MDSCs (*r*), and the MDSC death rate (*d*_*M*_). Since these can be identified using a sparse and noisy data set, we conclude that these are suitable parameters to vary during treatment personalization.

## 5. Experimental Data

We briefly describe the experimental data sets used in our mathematical modeling study. Optimization of treatment regimens and the occurrence of immune-related adverse events (irAEs) in simulated treated mice (Section 6) utilized three sets of murine data where anti-PD-1 and/or a CCR2 antagonist were administered.

In Tu et al. (2020), mice with implanted bladder or breast tumors were treated with anti-PD-1 and a CCR2 antagonist. Tumor volumes during treatment were measured for four treatment groups: no therapy, anti-PD1 monotherapy, CCR2 antagonist monotherapy, and combination therapy (Tu et al. (2020)[Figs. 2a and 4a]). Mice received 0.05 g of anti-PD-1 on days 14, 18, and 21 for bladder tumors and days 10, 14, and 17 for breast tumors post-implantation. Mice also received 2 mg/kg of the CCR2 antagonist daily from day 14 to 30 for bladder tumors and day 10 to 25 for breast tumors. This study used female C57BL/6J mice, which were 9 weeks old or older at the start of treatment. We assume that these mice weigh an average of 24 g (Gargiulo et al. (2014)). Thus, each dose amounts to roughly 0.05 mg of the CCR2 antagonist.

**Figure 2:**
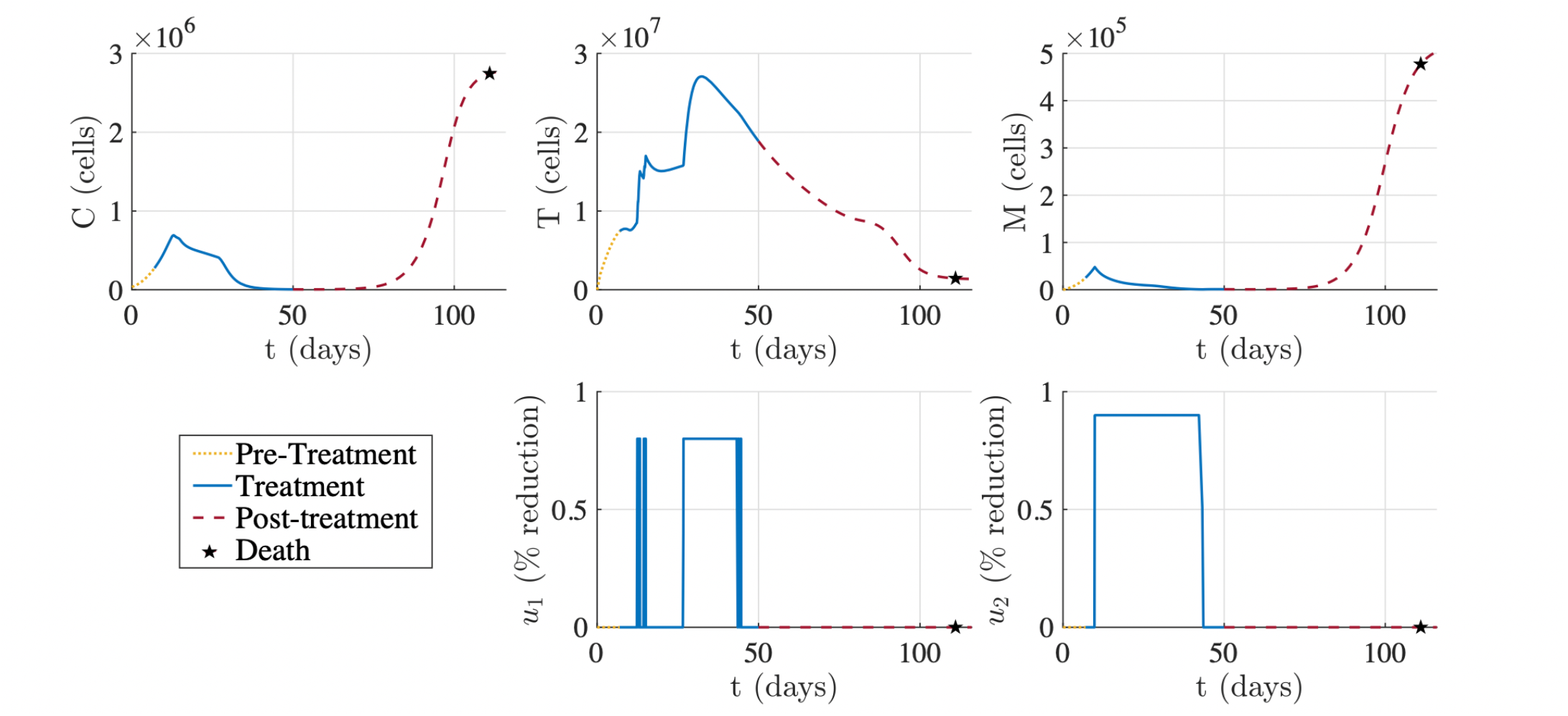
Example of optimized treatment for the best fitting parameter set from Table 1 according to the constraints stated in (6). Tumor was implanted on day 0. Treatment lasted from day 7 to day 50. Virtual subject surpassed tumor mortality threshold and thus died on day 111.

Flores-Toro et al. (2020) administered anti-PD-1 and a CCR2 antagonist to glioma-bearing mice and evaluated therapeutic efficacy. Starting on day 7 after glioma implantation, the CCR2 antagonist was administered twice daily for 21 days at a dose of 90 mg/kg. Anti-PD-1 administration also started on day 7 with a loading dose of 500 *μ*g/100 *μ*L, followed by 4 doses of 200 *μ*g/100 *μ*L every three days. Approximately 60% of mice implanted with KR158 glioma cells survived to 100 days post-implantation (Flores-Toro et al. (2020)[Fig. 4B]). In mice, irAEs are determined by the presence of immune infiltrates in organs outside the tumor (Ferreira et al. (2021); Adam et al. (2021)). Analysis of the blood and lymph nodes showed no change in immune infiltrates when mice were treated with or without the combination immunotherapy (Flores-Toro et al. (2020)[Fig. S6]). Therefore, we conclude that there were no irAEs reported due to the regimen itself.

Adam et al. (2021) evaluated the occurrence of irAEs in mice due to antiPD-1 treatment in combination with CFA boosters or anti-CTLA-4. In this experiment, mice were treated with anti-PD-1 twice a week for a maximum of 6 weeks, which is more than the regimen in Flores-Toro et al. (2020). irAEs occurred most frequently in the liver and lung tissue. These events could be related to anti-PD-1, anti-CTLA-4, and/or the CFA boosters.

## 6. Treatment Personalization

In this section, we personalize treatment regimens for virtual mice and then predict treatment outcomes. First, in Section 6.1, we estimate the rate at which anti-PD-1 and the CCR2 antagonist reduce inhibition by PD-L1PD-1 and recruitment of MDSCs, respectively, using data from Tu et al. (2020), and use this to determine bounds *b*_1_ and *b*_2_ in (4) for treatment. Then, in Section 6.2, we demonstrate an optimized treatment regimen for an average mouse according to the optimal control problem stated in Section 3. Lastly, in Sections 7.1–7.3, we obtain two sets of 10,000 virtual mice by randomly sampling the 5 practically identifiable parameters determined in Section 4. The first cohort represents subjects with a general tumor, and it is used to determine general markers for mortality and quality of life concerns during treatment and survival outcomes post-treatment in Sections 7.1 and 7.2, respectively. The second cohort represents subjects with GBM, and in Section 7.3, the median survival of the GBM cohort is evaluated and outcomes are compared between the two cohorts.

### 6.1. Therapeutic bounds for anti-PD-1 and CCR2 antagonist

We converted the tumor volumes reported in Tu et al. (2020) to cell counts by assuming that density of epithelial tumors is approximately 10^8^ cell/cm^3^ (Del Monte (2009)). Using the average tumor size at each time point, we re-estimated the five practically identifiable parameters (*λ*_*C*_, *η, ρ, r*, and *d*_*M*_) for the treatment-free groups and set all other parameters to the “Best Fit” values listed in Table 1. Given that the tumor sizes were an order of magnitude larger than the glioma data in Anderson et al. (2023), we also re-estimated the carrying capacity, *C*_max_. In each of the eight scenarios (2 without treatment, 6 with treatment), we obtained parameter distributions (Appendix E) by accepting 20,000 parameter sets of lowest error from a set of 100,000 samples using the approximate Bayesian computation (ABC) rejection method (Sunnåker et al. (2013); Liepe et al. (2014); Anderson et al. (2023)).

To quantify the efficacy of the doses reported in Tu et al. (2020) (see Section 5), we replace the inhibition/recruitment percent reductions *u*_1_ and *u*_2_ in model (1) with 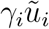, where *γ*_*i*_ (grams^−1^) is the efficacy of a therapeutic dose, 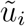 (grams), for *i* = 1, 2. Using the treatment-free parameters of best fit in Table E.3, we obtained estimates of *γ*_1_, *d*_1_, *γ*_2_, and *d*_2_ for model (1) and additional equations

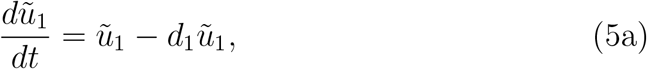

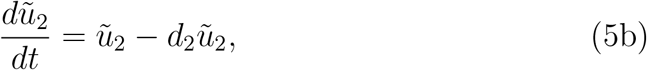

where *d*_1_ and *d*_2_ (day^−1^) are the rates of decay of anti-PD-1, 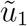, and the CCR2 antagonist, 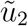, respectively. For the groups receiving treatment, parameter summary statistics are listed in Table E.4.

The “Best Fit” values in Table E.4 align with the mode of their respective distributions, each of which had distinctive peaks, resulting from the ABC rejection method. The only exception is *γ*_1_ for anti-PD-1 monotherapy in bladder cancer, which displayed a much higher mode of 1.80*×*10^4^ compared to the best fit of 6.21 *×*10^3^. However, this higher mode corresponds to all other *γ*_1_ values for best fit, mean, median, and mode listed in Table E.4.

To calculate estimates of the maximum percent reduction of the PD-L1PD-1 inhibition rate by anti-PD-1, *b*_1_, and the maximum percent reduction of the MDSC recruitment rate by the CCR2 antagonist, *b*_2_, as in (4) where 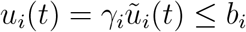, we observe that the biological mechanisms of anti-PD-1 and the CCR2 antagonist ensure that 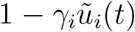 should be bounded below by 0 and above by 1. The largest bounds possible given the combination treatment group values for *γ*_1_ and *γ*_2_ (Table E.4) and doses from Tu et al. (2020) indicate that *b*_1_ = 0.8 and *b*_2_ = 0.9.

### 6.2. Optimized treatment regimen

We demonstrate an optimized treatment regimen for a single virtual mouse and compare it to previous experimental regimens described in Section 5. The optimal control problem (3) is solved using GPOPS-II (Patterson and Rao (2014)).

#### 6.2.1. Explicit problem statement

On day 0, the virtual mouse is implanted with 35,000 glioma cells (Appendix D). At that time, we assume that there is a small T cell population and no MDSCs at the tumor site, which corresponds to data in Anderson et al. (2023)[Fig. 3], giving the initial conditions *C*(0) = 35, 000, *T* (0) = 100, and *M* (0) = 0. Treatment starts at *t*_0_ = 7 days after implantation and continues at most until *t*_*f*_ = 50 days. The treatment start time corresponds to the regimen in Flores-Toro et al. (2020), and the maximum treatment duration was chosen in accordance with the Adam et al. (2021) anti-PD-1 toxicity study, which treated mice for a maximum of 6 weeks.

**Figure 3:**
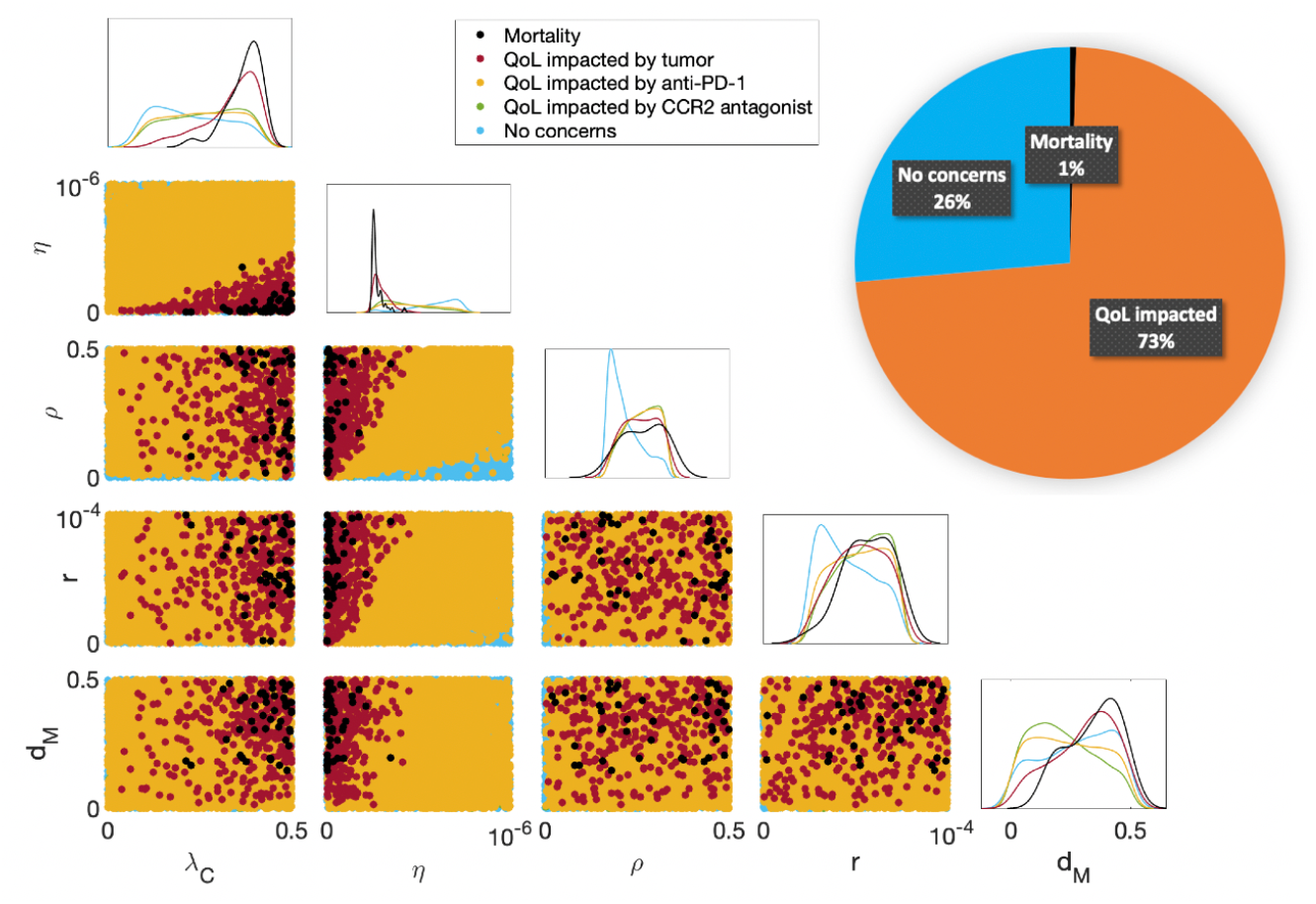
Mortality and quality of life (QoL) concerns during the treatment period of day 7 to day 50. The pie chart divides the cohort into the three categories of mortality, impacted QoL, and no concerns. Since most of the cohort experiences an impacted QoL, this shows that a more aggressive regimen is needed to control tumor sizes. The diagonal displays histograms for five different categories of mice according to each practically identifiable parameter from Section 4. The scatter plots beneath the diagonal show 2D projections of parameters and their resulting mortality and QoL concerns. A high tumor growth rate (*λ*_*C*_) and a low T cell kill rate (*η*) predict mortality during the treatment period. A high MDSC death rate (*d*_*M*_) was also a predictor of mortality during treatment, while a low MDSC death rate was associated with QoL concerns due to the CCR2 antagonist. The best predictors of no concerns during treatment were low T cell inhibition rates by PDL1-PD-1 (*ρ*) and by MDSCs (*r*).

Thus, the optimal control problem (3) becomes

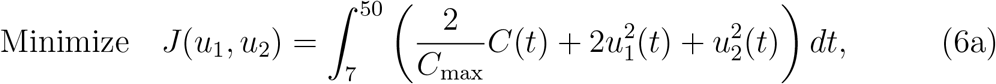

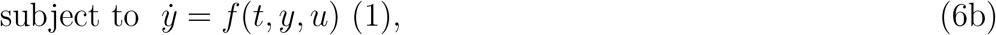

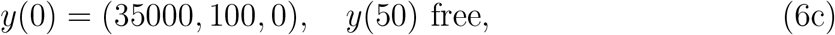

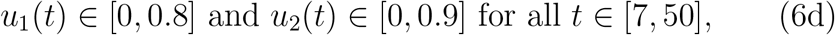

where *C*_max_ is the tumor carrying capacity. Weights for the tumor burden (*C*) and anti-PD-1 (*u*_1_) were twice as much as the CCR2 antagonist (*u*_2_), since the lower toxicity profile of the CCR2 antagonist indicates that it is more essential to minimize the tumor and anti-PD-1. Different integer combinations from 1 to 10 were tested for the three weights, and weight combinations where the tumor burden and anti-PD-1 were weighted greater than the CCR2 antagonist yield similar results.

We assume that the cumulative percent reduction by anti-PD-1 and the CCR2 antagonist satisfy

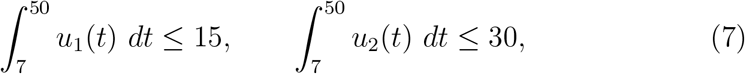

as this roughly aligns with the maximum number of treatment days in the experimental regimens described in Section 5. Lastly, to ensure the treatment terminates if it cannot decrease the tumor size, we assume that the final tumor state is bounded above by the tumor size at the start of treatment, *C*(7). Implementing these conditions, we solve the problem (6) with GPOPS-II optimization software (Patterson and Rao (2014)). GPOPS-II utilizes Gaussian quadrature collocation methods to approximate the continuous optimal control problem as a sparse nonlinear programming (NLP) problem. This NLP problem is then solved to determine the optimal treatment regimen (*u*_1_(*t*), *u*_2_(*t*)) to minimize the objective functional (6a).

#### 6.2.2. Optimized regimen for an average mouse

Figure 2 displays the optimized treatment regimen for the simulated mouse associated with the “Best Fit” parameter set from Table 1. The optimized treatment results in a substantial decrease in tumor burden (*C*) where the final tumor size at the end of treatment is 1.9% of the initial tumor size on day 7. Anti-PD-1 (*u*_1_) is administered over four intervals, each of which are at the maximum allowable percent reduction. The first two anti-PD-1 doses are at days 12 and 14, and then anti-PD-1 is maintained at a constant level from day 26 to 44, where it is then released quickly and then given one last dose. Comparably, the CCR2 antagonist (*u*_2_) is administered at its maximum allowable percent reduction from day 10 to 42 and then discontinued. Achieving the maximum allowable percent reduction directly correlates with concentrating the therapy at the glioma site when treated at its maximum effective dose, which is defined as the dose “beyond which additional benefit would be unlikely to occur” (Guideline (1999)). Therefore, Figure 2 showcases treatment with anti-PD-1 and the CCR2 antagonist at their respective maximum effective dose.

Depending on the decay rates of anti-PD-1 and the CCR2 antagonist, maintaining these drugs at a constant percent reduction does not necessarily mean constant administration. Flores-Toro et al. (2020) administered antiPD-1 every 3 days and the CCR2 antagonist twice daily, which was very similar to the regimen in Tu et al. (2020). Assuming that this treatment frequency produces a roughly constant concentration of each drug at the glioma site, Figure 2 suggests that an optimal treatment for glioma-bearing mice is anti-PD-1 administration on day 12 and then every three days starting on day 26 for six additional doses and CCR2 antagonist administration twice daily from day 10 to day 42. Both therapies are to be administered at their maximum effective doses for each administration.

#### 6.2.3. Comparison to experimental treatment regimens

Compared to the experimental regimens with combination immunotherapy in Flores-Toro et al. (2020) and Tu et al. (2020), the computed optimal treatment regimen starts later after tumor implantation but lasts longer. In Flores-Toro et al. (2020), treatment started on day 7 after glioma implantation for both therapies, while our optimized regimen suggests initiation of the CCR2 antagonist on day 10 and anti-PD-1 on day 12. These start times are more similar to the regimen for breast tumors in Tu et al. (2020), which started the CCR2 antagonist on day 10 and anti-PD-1 on day 17.

Further, Flores-Toro et al. (2020) had 5 doses of anti-PD-1 and 21 days of CCR2 antagonist treatment compared to our increased 7 doses of anti-PD-1 and 32 days of the CCR2 antagonist. While this is more anti-PD-1 doses than either Flores-Toro et al. (2020) or Tu et al. (2020), it is less than the maximum 12 anti-PD-1 doses for mice tested in Adam et al. (2021). We also see an increase in the number of CCR2 antagonist treatment days compared to Tu et al. (2020), which treated tumors with a CCR2 antagonist for a maximum of 26 days.

The most notable difference between our computed optimized regimen and the experimental regiment in Flores-Toro et al. (2020) was the initial dose of anti-PD-1. While Flores-Toro et al. (2020) had a higher loading dose with lower subsequent maintenance doses, our regimen suggests that anti-PD-1 should be administered at the same dose. Further, our regimen suggests the first dose is two weeks before subsequent anti-PD-1 doses, which is different from the constant dose frequency of every 3 days in Flores-Toro et al. (2020).

## 7. Mortality and Morbidity Analysis in a Virtual Cohort

We predict the occurrence of mortality and impacted quality of life (QoL) during the treatment interval in a virtual murine cohort. We set a variety of thresholds to categorize the cohort and consider dose escalation for more aggressive tumors (Section 7.1). Then, we evaluate the long-term outcomes of mice in response to therapy (Section 7.2) and repeat the mortality and morbidity analyses with a GBM-specific virtual murine cohort (Section 7.3). We assume that therapeutic efficacy is determined by tumor reduction and QoL. Since impacted QoL can range from mild to severe, more emphasis is placed on the role of tumor reduction for treatment efficacy.

### 7.1. During treatment: Mortality and Quality of Life (QoL) concerns

Factors that affect mortality include the tumor size in relation to its carrying capacity (*C*_max_), and factors affecting QoL include tumor burden and adverse events due to drug toxicities.

We set QoL thresholds for anti-PD-1 and the CCR2 antagonist based on the regimens of Adam et al. (2021) and Flores-Toro et al. (2020) as described in Section 5. We assume that QoL is not impacted unless anti-PD-1 or the CCR2 antagonist is administered for a large number of days at approximately their maximum percent reduction. Additionally, since toxicity could be related to the cumulative dose instead of the number of doses, we bound the cumulative percent reduction of the two drugs.

Letting *C* be the number of tumor cells at any time from day 0 to 50, we therefore define the following mortality and QoL thresholds:

- *Mortality:*

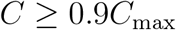

- *QoL impacted by the tumor:*

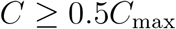

or *C* ≥ *C*(0) = 35, 000 for more than half the treatment period

- *QoL impacted by anti-PD-1:*

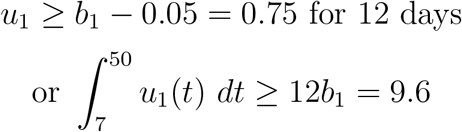

- *QoL impacted by CCR2 antagonist:*

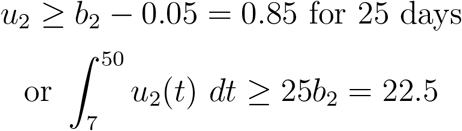

If a subject does not surpass any of the mortality or QoL thresholds, we assume that there are no concerns during the treatment period.

#### 7.1.1. Categorization of virtual cohort

10,000 virtual subjects were generated by approximate uniform sampling of the five practically identifiable parameters (Section 4) using Latin hypercube sampling within the parameter ranges from Anderson et al. (2023). All other parameters were fixed to their “Best Fit” values in Table 1. The optimized treatment regimen is determined for each virtual mouse as described in Section 6.2. In Figure 3, we plot 2D projections of the 10,000 virtual subjects according to their mortality and QoL concerns during the treatment period. Along the diagonal, there are 1D projected histograms of the mortality and QoL concerns according to each parameter. A pie chart divides the cohort into three separate categories: mortality, impacted QoL, and no concerns.

Of the 10,000 virtual subjects, 73% experience an impacted QoL, largely due to anti-PD-1 (99.6% of subjects with concerns). The CCR2 antagonist is less likely than anti-PD-1 to impact QoL, as 59.3% of the concerns are related to this therapeutic. This suggests that most subjects experience some level of side effects due to treatment administration. Since more subjects experience side effects due to anti-PD-1, this is likely due to the lower QoL threshold and the anti-tumor benefit of exceeding this threshold. Approximately 4% of the 10,000 mice experience mortality (1%) or an impacted QoL due to the tumor size (3%), which shows that optimized regimens can control the tumor well during treatment.

Figure 3 indicates that subjects who die during treatment have a high tumor growth rate (*λ*_*C*_), low T cell kill rate (*η*), and a high MDSC death rate (*d*_*M*_). In these cases, the mouse’s tumor is too aggressive and they are too immunocompromised to respond to therapy. Although a high MDSC death rate (*d*_*M*_) is associated with death, it is also associated with having neither mortality nor QoL concerns. Therefore, it is a poor biomarker for outcomes during the treatment period. On the other hand, a low MDSC death rate indicates that the subject may experience QoL issues due to the CCR2 antagonist. For these mice, their MDSC population is living longer, requiring an increase in CCR2 antagonist administration to control the MDSC-induced immune suppression. Lastly, subjects presenting no mortality or QoL concerns had low inhibition rates by PD-L1-PD-1 (*ρ*) and by MDSCs (*r*). These mice experience less immune suppression from these inhibition mechanisms, and thus needed less of either immunotherapy.

Figure 4 illustrates the cumulative percent reduction of anti-PD-1 versus the CCR2 antagonist, which correlates with the cumulative dose (Section 6.1). As tumor aggression increases, anti-PD-1 and the CCR2 antagonist are increased linearly until reaching the upper bound for the cumulative percent reduction of anti-PD-1 specified in Section 6.2. Then, the CCR2 antagonist is increased until reaching its specified upper bound. In general, virtual subjects who died were given the highest allowed levels of both drugs, but this was unable to control their aggressive tumor growth. A handful of mice who died were treated at lower levels; this is likely due to even higher levels of tumor aggression, as increased tumor growth caused the treatment to terminate early.

**Figure 4:**
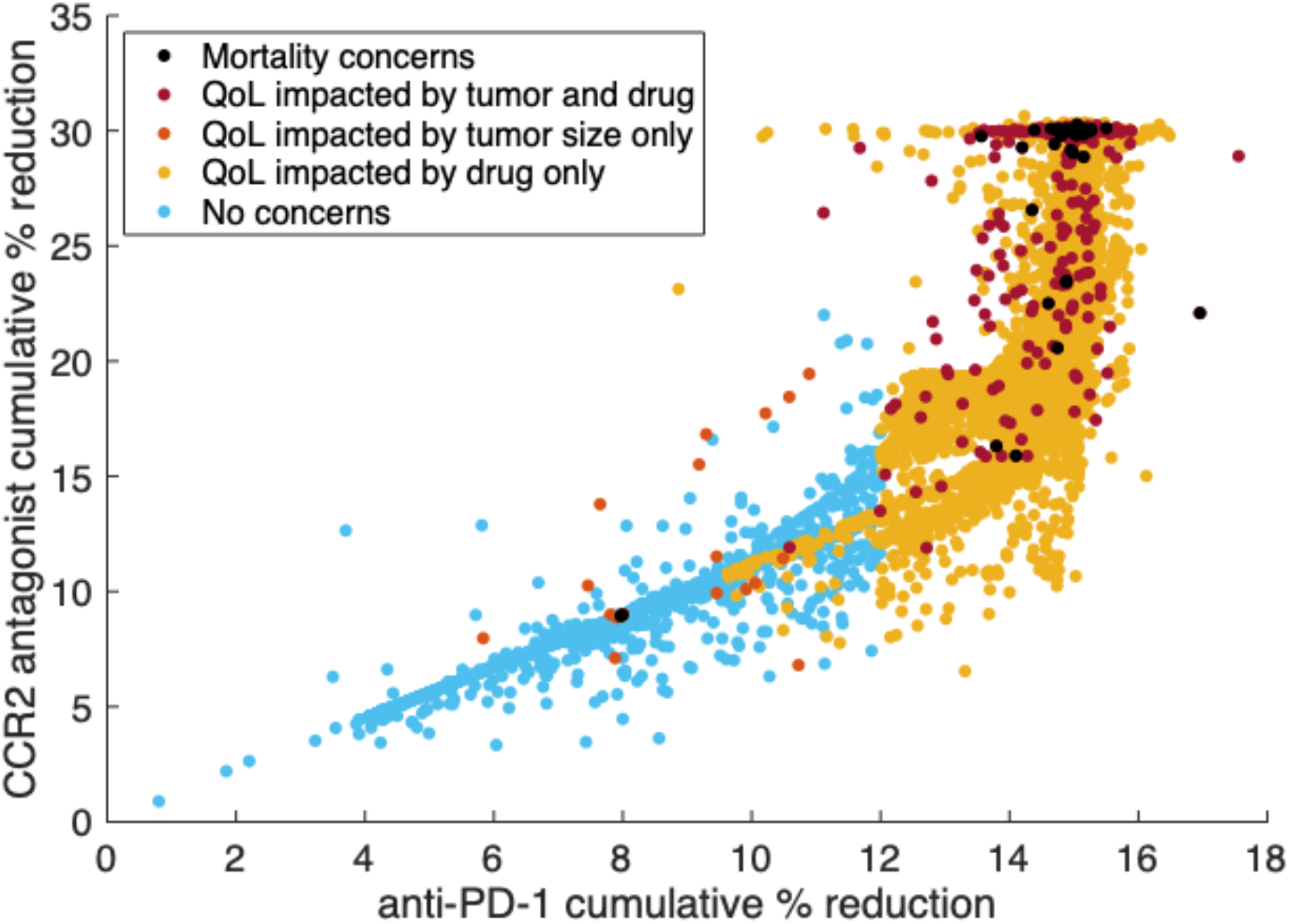
Scatter plot of the cumulative percent reduction of anti-PD-1 versus the CCR2 antagonist for each personalized treatment regimen from the virtual murine cohort. Subjects are categorized with respect to mortality and quality of life (QoL) thresholds from Section 7.1. For more aggressive tumors, the cumulative percent reductions of anti-PD-1 and the CCR2 antagonist increase linearly as needed until reaching the specified bound for anti-PD-1. Then, the CCR2 antagonist is increased until reaching its bound.

### 7.2. After treatment: Disease and progression-free survival, recurrence, and failure

Since it is often standard to evaluate 2-year and 5-year survival outcomes for human patients, we do the same but in terms of the mouse lifespan. The average lifespan of mice is 836 days (Kunstyr and Leuenberger (1975)). This value is based on data from C57BL/6 mice, which is the breed used in FloresToro et al. (2020) and Tu et al. (2020). Currently, in the US, the average lifespan is 76.1 years (Arias et al. (2022)), so we take a mouse “year” to be 11 days. Therefore, we evaluate survival outcomes after 22 and 55 days.

Similar to Section 7.1, we assume that a virtual subject dies if their tumor nears its carrying capacity (*C*_max_) post-treatment. Assuming a tumor is smaller than the mortality threshold (i.e., the subject is alive), a subject is categorized as progression-free if their tumor is decreasing after treatment or if there is little variation in the tumor sizes on the last 10 days of the survival time period. Subjects with less than 1 tumor cell remaining are considered disease-free.

Let 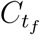 be the tumor cell count on the final day of treatment, and let *C*_*n*_ be the number of tumor cells *n* days after treatment. We define the following thresholds, evaluated at *n* = 22, 55:

- *Death:*

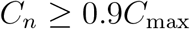

- *Progression-free survival:*

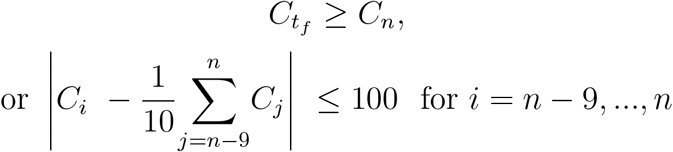

- *Disease-free survival:*

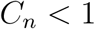

We assume that all subjects not in these three categories experience tumor recurrence.

#### 7.2.1. Survival analysis

Figure 5 illustrates the results for 2-year and 5-year survival analysis. High tumor growth rate (*λ*_*C*_), low T cell kill rate (*η*), and high inhibition rates by PD-L1-PD-1 (*ρ*) and by MDSCs (*r*) are all indicators of death by both year 2 and year 5. Mice with these tumors are immune-compromised with more aggressive tumors. A low MDSC death rate (*d*_*M*_) is a marker for tumor recurrence by year 2 and death by year 5, while a high *d*_*M*_ is a marker of disease-free survival by year 5. Subjects with a high *d*_*M*_ have a mechanism by which to internally control MDSC-induced immune suppression, while subjects with a low *d*_*M*_ are gradually overcome by this form of immune suppression, leading to tumor recurrence and death. A low *λ*_*C*_ is also a marker of disease-free survival by year 5, as the immune response of these subjects can more easily control the slower tumor growth.

**Figure 5:**
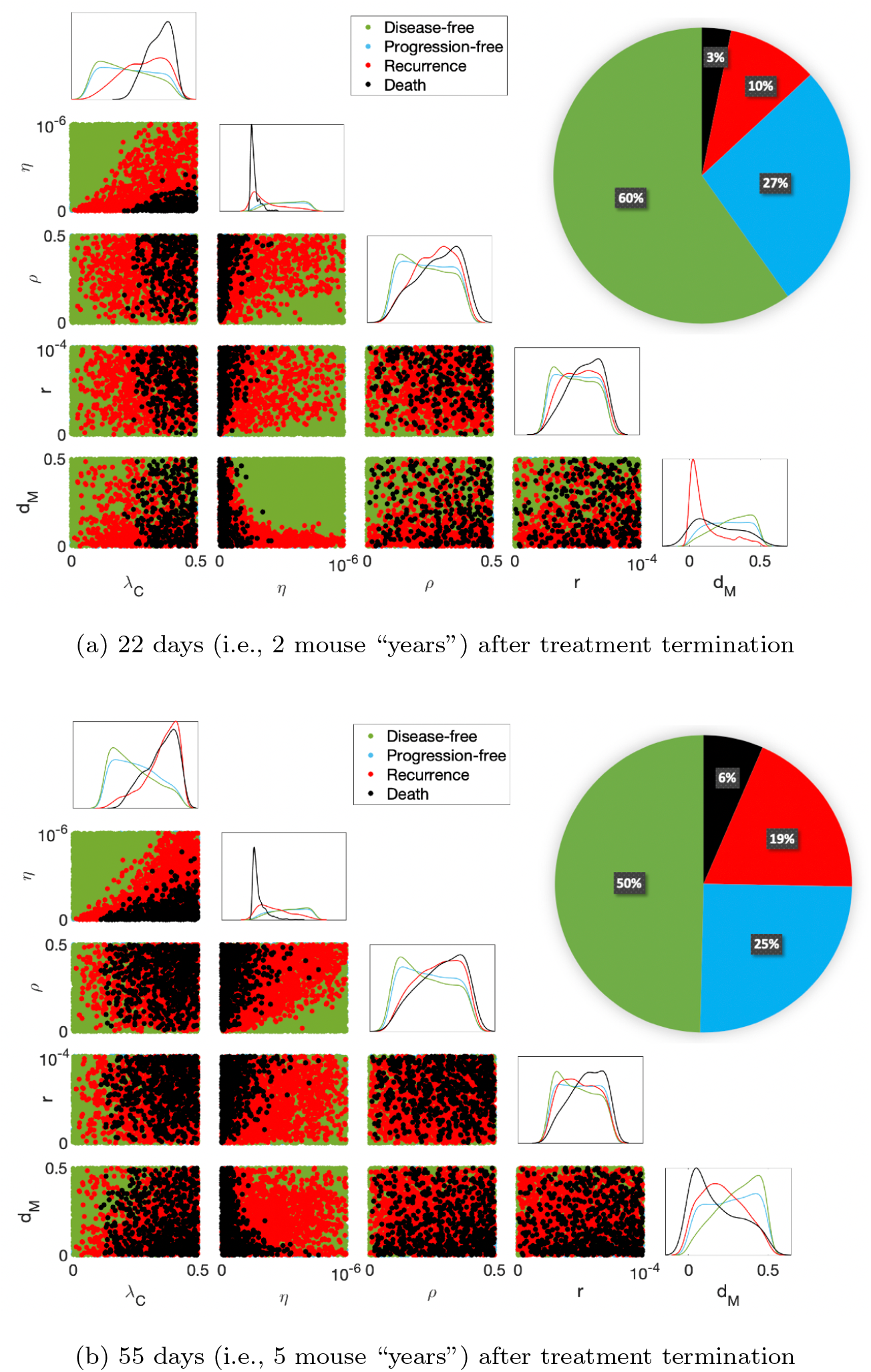
Survival analysis post-treatment. The pie chart divides 10,000 subjects into the 4 survival categories. The diagonal displays a histogram for each category according to each practically identifiable parameter from Section 4. The scatter plots beneath the diagonal show 2D projections of parameters and predicted disease-free survival, progression-free survival, tumor recurrence, and death. A high tumor growth rate (*λ*_*C*_), low T cell kill rate (*η*), high inhibition rates by PD-L1-PD-1 (*ρ*) and by MDSCs (*r*), and a low MDSC death rate (*d*_*M*_) are markers of treatment failure over time.

### 7.3. Glioblastoma (GBM)-specific virtual murine cohort

The general tumor virtual murine cohort studied in Sections 7.1–7.2 was sampled approximately uniformly from the identifiable parameter space in order to clearly identify qualitative trends. However, to obtain more realistic quantitative predictions, we generate a GBM virtual murine cohort with parameter values sampled according to GBM-specific probability distributions in Anderson et al. (2023)[Table 2]. In particular, each practically identifiable parameter (*λ*_*C*_, *η, ρ, r*, and *d*_*M*_) is randomly sampled according to its probability distribution obtained from GBM murine data, while the remaining parameters are set to the “Best Fit” value from Table 1.

Figure 6a categorizes the virtual GBM cohort according to quality of life (QoL) and survival outcomes during and after treatment. Compared to pie charts in Figures 3 and 5, the GBM cohort is more likely to receive a more aggressive personalized treatment regimen, as seen by an increase in QoL concerns (73% compared to 82%). Further, virtual GBM subjects are more likely to experience death or tumor recurrence after treatment conclusion. For instance, 55 day survival outcomes show that mortality increases from 6% to 15% and tumor recurrence increases from 19% to 34%.

**Figure 6:**
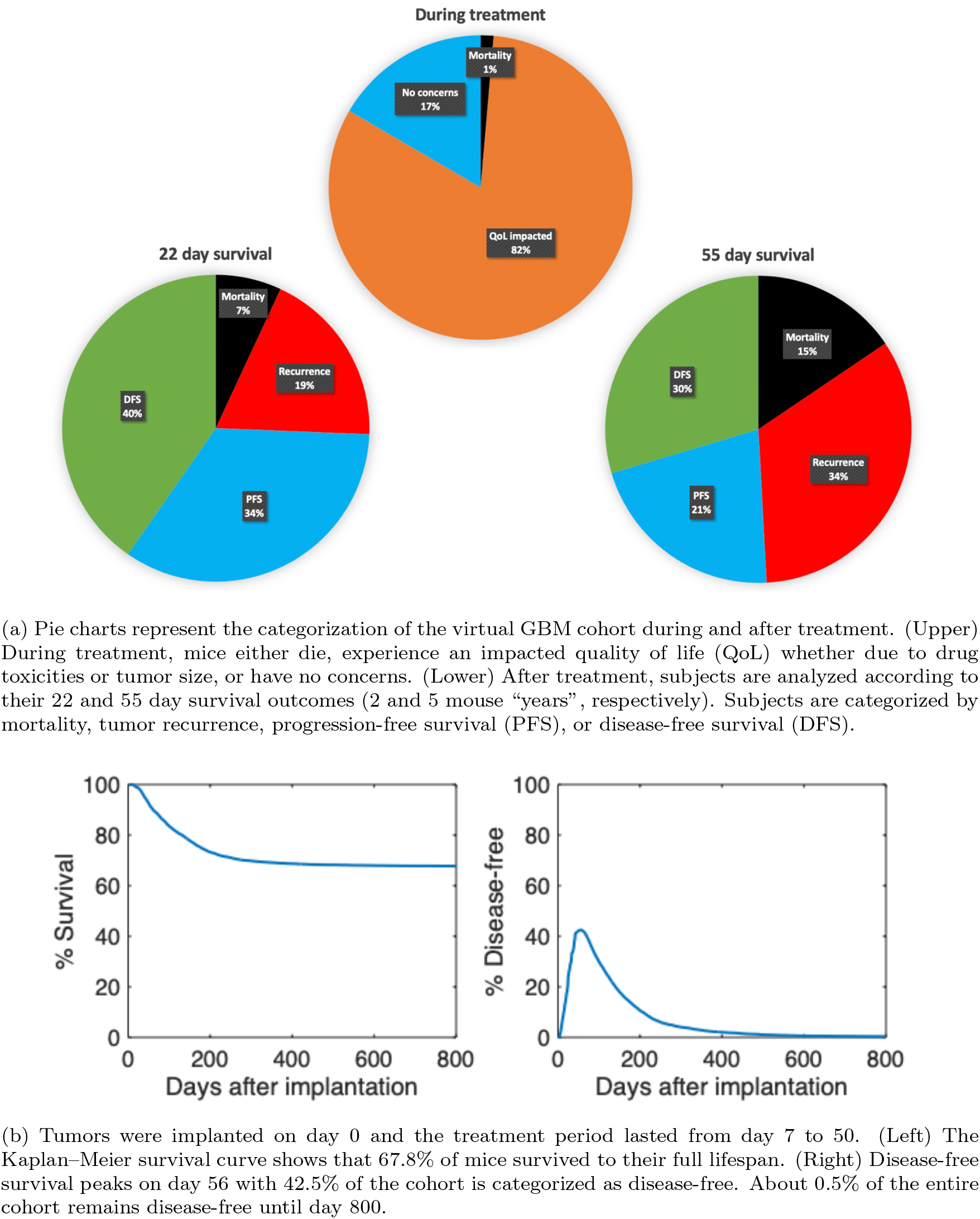
Survival analysis for the cohort of 10,000 virtual GBM mice after receiving a personalized optimal treatment regimen with anti-PD-1 and the CCR2 antagonist. The virtual cohort was sampled using the GBM-specific distributions of the 5 practically identifiable parameters from Anderson et al. (2023)[Table 2].

Figure 6b shows the Kaplan–Meier survival curve for the virtual GBM cohort and the percent of subjects that are disease-free at any time. We assume that tumor implantation on day 0 occurs at least several weeks after birth, so the remaining mouse lifespan is represented in the figure. By day 300, the Kaplan–Meier survival curve starts to plateau until 67.8% of subjects remain at day 800. The percent of disease-free subjects reaches 42.1% at the end of the treatment period (day 50) and then peaks six days later at 42.5%. This continues to decline until day 800 when 0.5% of the cohort remains disease-free.

## 8. Discussion

Glioblastoma (GBM) is a highly aggressive primary brain tumor in need of improved treatment strategies. A combination immunotherapy regimen with anti-PD-1 and a CCR2 antagonist showed efficacy in preclinical murine models (Flores-Toro et al. (2020)). In this paper, we extend the GBMimmune dynamics model from Anderson et al. (2023) to include treatment with the combination immunotherapy and formulate a treatment optimization problem in terms of optimal control theory. The aim of this study was to obtain optimized, personalized treatment regimens for virtual subjects and predict markers of treatment success and failure.

Before optimization, parameter identifiability analysis (Section 4) was conducted with the treatment-free model to determine parameters to highlight during personalization. The model was found to be structurally identifiable with respect to tumor cell count data, meaning that given enough noise-free tumor data, all model parameters can theoretically be identified. Practical identifiability analysis was then performed, and results show that murine data on tumor, T cells, and MDSCs over the course of six time points was able to identify five parameters, namely the tumor growth rate (*λ*_*C*_), the T cell kill rate (*η*), the inhibition rates by PD-L1-PD-1 (*ρ*) and by MDSCs (*r*), and the MDSC death rate (*d*_*M*_). Thus, since these parameters can be identified despite sparse and noisy data, treatment personalization results regarding these 5 parameters can be used in practice.

Within the subspace of the five practically identifiable parameters, we sampled 10,000 virtual murine subjects and then optimized the combination immunotherapy for each subject. Mice were then categorized according to their predicted survival, mortality, and quality of life outcomes (Sections 7.1 and 7.2), which led to identifying markers for treatment failure and success. We specified thresholds to categorize each subject, however, thresholds are more arbitrary than our work demonstrates, so results are more suited to identify population trends rather than concretely categorize an individual subject.

As can be expected, subjects with high tumor growth rates (*λ*_*C*_) and low T cell kill rates (*η*) were more likely to die during and after treatment as seen in Figures 3 and 5. For these mice, it is impossible to amplify the immune response enough with the combination immunotherapy to overcome the aggressive tumor growth. Subjects with higher inhibition rates by PD-L1-PD-1 (*ρ*) and by MDSCs (*r*) require more aggressive treatment with anti-PD-1 and the CCR2 antagonist, respectively, and are more likely to die after treatment due to the increased immune suppression. Unexpectedly, the MDSC death rate (*d*_*M*_) was a better predictor for a more aggressive treatment with the CCR2 antagonist than *r*. Further, *d*_*M*_ was also a better indicator of longterm survival than either of the immune suppression parameters *ρ* or *r*. A low *d*_*M*_ predicts tumor recurrence by “year” 2 (day 22 post-treatment in mice) and death by “year 5” (day 55). The 5-year survival for GBM has remained largely unchanged despite improvements to the median and short-term overall survival in recent years (Cantrell et al. (2019); Siegel et al. (2023)); therefore, obtaining *d*_*M*_ as a predictor of treatment success and failure by year 5 is a useful step forward. Note that the GBM-specific distributions of the 5 identifiable parameters in Anderson et al. (2023) were all right-skewed except for the MDSC death rate, which exhibited a normal distribution. Thus, for GBM specifically, this decreases the importance of trends identified on the upper ranges for the tumor growth rate, T cell kill rate, and the inhibition rates by PD-L1-PD-1 and by MDSCs, and increases the importance of the MDSC death rate as a marker of long-term treatment failure or success.

Given data limitations, we bounded the anti-PD-1 and CCR2 antagonist treatments by an estimate of the maximum percent that they reduce PDL1-PD-1 inhibition and MDSC recruitment, respectively. This maximum percent reduction corresponds to the drug concentration at the tumor site when the drug is administered at its maximum effective dose. As we used time course data from Tu et al. (2020), which studied bladder and breast tumor response to combination immunotherapy, we would be able to better predict the maximum percent reduction and drug decay rates for our GBM-specific model with data from gliomas. In the future, we seek to obtain this data, re-estimate the efficacy and decay of anti-PD-1 (*γ*_1_, *d*_1_) and CCR2 antagonist (*γ*_2_, *d*_2_) in glioblastoma, and predict accurate doses and frequencies.

With these constraints in mind, we optimized treatment for a virtual GBM murine subject represented by the parameter set of “Best Fit” from Table 1 in Section 6.2. This parameter set was obtained in Anderson et al. (2023) using average data from 17 glioma-bearing mice across six time points. Therefore, although the regimen in Figure 2 is optimized for a specific mouse, it represents a suitable regimen for the average mouse with GBM.

Figure 2 suggests that the optimal treatment for glioma-bearing mice is an increased tumor site concentration of anti-PD-1 on days 12, 14, and 26 to 44, and the CCR2 antagonist from days 10 to 42, where both drugs are administered at a dose and frequency that allows them to obtain their maximum percent reductions during these periods. Compared to the murine regimen for GBM in Flores-Toro et al. (2020), the computed optimized treatment starts later after tumor implantation but lasts longer. Both regimens had a single prolonged period of treatment with the CCR2 antagonist. Unlike the computed optimized regimen, the doses of anti-PD-1 were evenly spaced in Flores-Toro et al. (2020). Further, Flores-Toro et al. (2020) had a higher loading dose compared to subsequent maintenance doses, but the optimized regimen suggests anti-PD-1 be treated at a constant level.

Figure 2 also shows the system dynamics of the virtual GBM mouse posttreatment. According to the survival thresholds in Section 7.2, we have tumor recurrence by “year” 2 post-treatment (day 72). By “year” 5 (day 105), the subject is nearing death and by day 111, the mortality threshold is exceeded and the subject dies. Although death still occurred, the survival time of this virtual subject increased by 79 days, since the mortality threshold is exceeded by day 32 without treatment. Our predicted survival time for the average non-treated mouse is supported by the experimental median survival of 28 days for non-treated glioma bearing mice in Flores-Toro et al. (2020)[Fig. 4B]. Further, although Flores-Toro et al. (2020) only evaluated survival until 100 days post-implantation, the percent survival at day 100 is approximately 60%, thus corresponding to the 111 day predicted survival of the average mouse post-implantation and treatment with the combination immunotherapy. Additionally, the Kaplan–Meier survival curve for the virtual GBM cohort in Figure 6b illustrates an 84% survival on day 100, showing that the personalized optimal regimens can improve outcomes in GBM compared to the current experimental regimen.

While this work was specifically applied to GBM, similar parameter ranges in Figures 3 and 5 could be used for other cancers. Therefore, the conclusions regarding markers for survival, mortality, and quality of life apply to all cancers being treated with anti-PD-1 in combination with a CCR2 antagonist. Results from our GBM subject in Section 6.2 and GBM-specific cohort in Section 7.3, however, do not apply to other cancers. In order to replicate these findings for other cancers, it would be necessary to re-estimate practically identifiable parameters and their probability distributions to represent a particular tumor type and then optimize therapy. This further exploration could be useful for cancers like pancreatic, bladder, and breast cancer, which have all been treated concurrently with anti-PD-1 and a CCR2 antagonist (Orth et al. (2019); Tu et al. (2020)). Since tumor size can vary significantly depending on location, it will be necessary to also re-estimate the tumor carrying capacity (*C*_max_) for different cancers. However, *C*_max_ is practically unidentifiable, so the assumption will need to be made that a fairly realistic prediction of this parameter is able to be obtained. Further, since our objective functional uses *C*_max_ as a weight to balance minimizing the tumor burden and drug toxicities, one should consider fixing the *C*_max_ estimate for an entire tumor type to maintain consistency in treatment predictions for a cohort. Although it was necessary to use this practically unidentifiable parameter to weight the objective functional, this aspect of our approach partially limits personalization of the optimal control method.

Data limitations make it difficult to contain all the complexities of GBM within any single model. Our model, as well as the foundational Anderson et al. (2023) model, differ from previous mathematical representations of GBM by including myeloid-derived suppressor cells (MDSCs). However, GBM exhibits a particularly complex immune environment with other immune cells, molecules, and checkpoints (Duerinck et al. (2023)). Like our GBM-immune model, Storey et al. (2020) included the PD-L1-PD-1 immune checkpoint while also modeling innate and adaptive immunity. In the GBMimmune model by Khajanchi and Nieto (2021), they included macrophages and two cytokines, namely TGF-*β* and IFN-*γ*. Santurio and Barros (2022) addressed CAR-T cells as well as the wider brain microenvironment by introducing neurons and glial cells. In the future, layering in additional layers of the immune and brain microenvironment as well as barriers to treatment, such as GBM’s highly heterogeneous nature and the blood-brain barrier (Cruz et al. (2022)), would enable the determination of more accurate responses to treatment.

In conclusion, our work extends the original GBM-immune dynamics model by Anderson et al. (2023) to include combination immunotherapy with anti-PD-1 and a CCR2 antagonist. The methods used in this paper can easily be implemented to improve therapeutics for other cancers. Optimized treatment regimens show an increase in survival compared to experimental regimens, and results also identify the MDSC death rate as a useful predictor of long-term survival for GBM patients. While our predicted regimens are specifically for mice, this work gives a basis for predicted treatment efficacy in humans. In the future, more work would need to be done to test the optimized regimens in mice and extend results to the human condition.

## CRediT authorship contribution statement

**Hannah G. Anderson:** Conceptualization, Methodology, Formal analysis, Software, Validation, and Visualization, Writing - original draft, Writing - review and editing. **Gregory P. Takacs:** Conceptualization, Methodology, Writing - review and editing. **Jeffrey K. Harrison:** Conceptualization, Methodology, Resources, Writing - review and editing. **Libin Rong:** Conceptualization, Methodology, Supervision, Writing - review and editing. **Tracy L. Stepien:** Conceptualization, Methodology, Supervision, Writing - review and editing.

## Code availability statement

The source code used to generate the results for this article is available through GitHub at https://github.com/stepien-lab/glioma-Tcell-MDSC-treatment [v1.0.0]. The code is platform independent and written in MATLAB.

## Acknowledgments

The authors acknowledge University of Florida Research Computing for providing computational resources and support that have contributed to the research results reported in this publication.

## Appendix A. CCR2 Antagonist Derivation

Myeloid-derived suppressor cells (MDSCs) are recruited to a tumor site chemokinetically via the CCL2-CCR2 and CCL7-CCR2 axes (Takacs et al. (2022)). Treatment with a CCR2 antagonist, 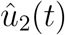, prevents this recruitment through blocking the CCR2 receptors of MDSCs. We model the net change of CCR2 receptors outside of the brain to be

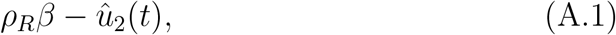

where *ρ*_*R*_ is the CCR2 expression of a single MDSC and *β* is the MDSC growth rate.

The rate of change of chemokines is expressed as

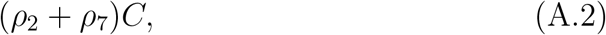

where *ρ*_2_ and *ρ*_7_ represent CCL2 and CCL7 expression, respectively, of tumor cells.

For simplicity, we assume that CCL2 and CCL7 exhibit the same rate of association, *a*, and dissociation, *d*, to the CCR2 receptor. Therefore, the chemical reaction can be viewed as

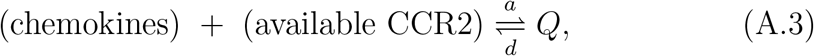

where *Q* represents the chemokine-CCR2 complex. Assuming that this reaction is in a quasi-steady state,

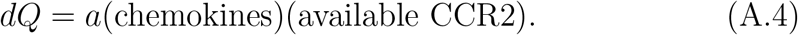

We conclude that the recruitment rate of MDSCs to the tumor site is

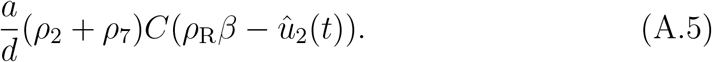

Due to parameter non-identifiability, we simplify this to

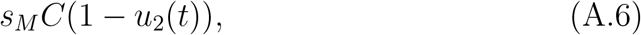

where

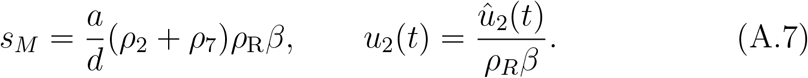

## Appendix B. Necessary Conditions for Optimality

The optimal control problem (3) can be reduced to minimizing the corresponding Hamiltonian with respect to *u*_1_ and *u*_2_,

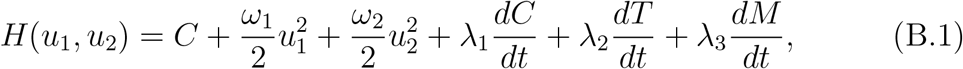

where *λ*_*i*_, *i* = 1, 2, 3, represents the adjoint function for the *i*th variable. The Hamiltonian is constructed to satisfy the first of several conditions for Pontryagin’s minimum principle (Chiang (1999)[Ch. 7]), namely 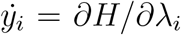, where *y*_*i*_ is the *i*th state variable.

Pontryagin’s principle also imposes two conditions on the adjoint functions,

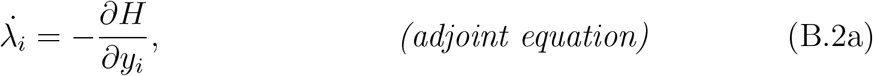

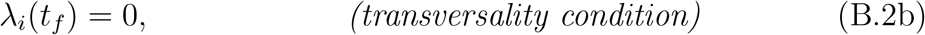

for *i* = 1, 2, 3, and where *t*_*f*_ is the final time point. Rewriting these conditions by substituting the Hamiltonian (B.1) and the system of model equations (1), we obtain

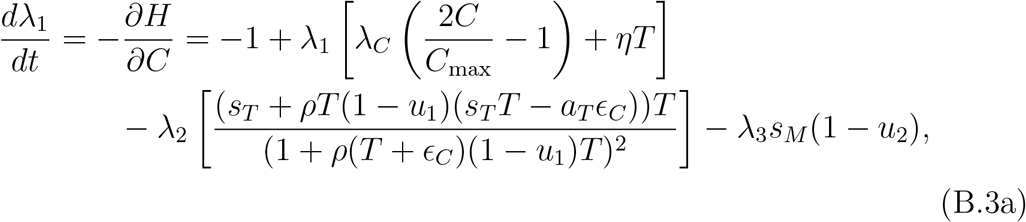

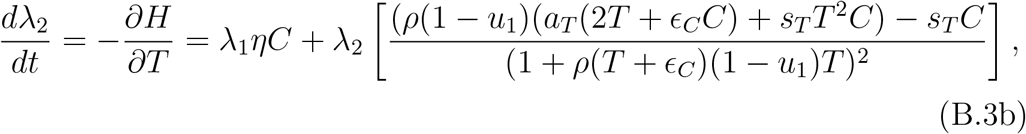

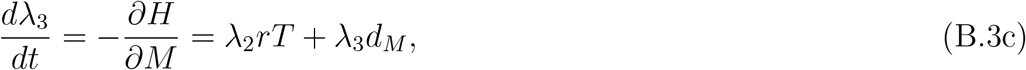

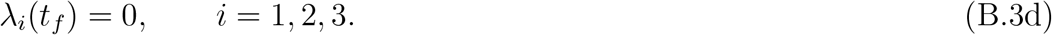

Now, we seek to characterize the optimal treatment regimen, (*u*_1_, *u*_2_).

### Theorem 1.

*There exists a unique optimal pair*, 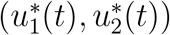, *that minimizes the Hamiltonian* (B.1), *characterized by*

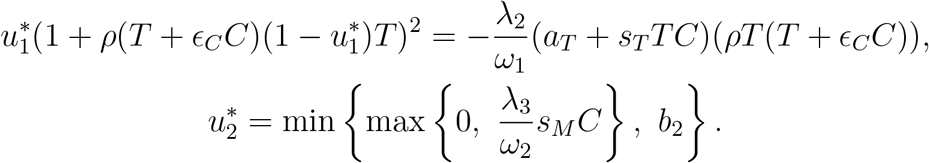

*Further, the characterization of* 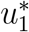 *is unique unless* 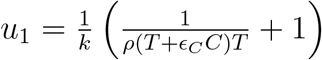 *where k* = 1 *or k* = 3.

*Proof*. Using the conditions of Corollary 4.1 in Fleming and Rishel (2012), we prove that there exists an optimal pair minimizing *J*(*u*_1_, *u*_2_) (3a) on 𝒰 (4).

𝒰 is closed and convex, and the integrand of the objective functional (3a),

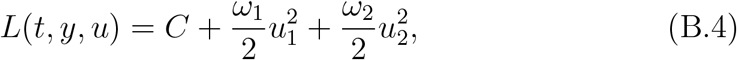

is convex with respect to *u*_1_ and *u*_2_. Further, 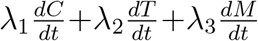 is continuous on a compact set since the state equations are continuous and bounded by Anderson et al. (2023). Lastly, Corollary 4.1 (Fleming and Rishel (2012)) requires that

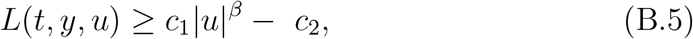

where *c*_1_ *>* 0 and *β >* 1. Since *C* ≥ 0 (Anderson et al. (2023)), this lower bound for *L*(*t, y, u*) is trivially fulfilled by 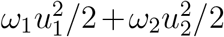. Thus, an optimal pair, (*u*_1_, *u*_2_), exists that minimizes the Hamiltonian (B.1).

When we minimize the Hamiltonian (B.1) with respect to *u*_1_, we obtain an implicit characterization for *u*_1_,

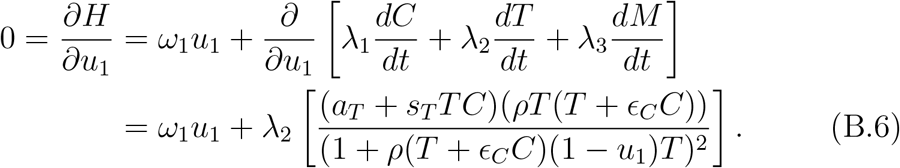

This characterization is unique under certain conditions. To exhibit this, we first rearrange (B.6) as

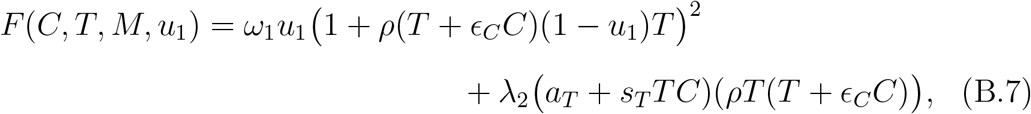

and differentiate with respect to *u*_1_,

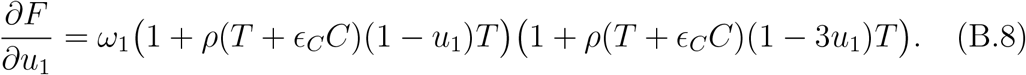

By the implicit function theorem, if ∂*F/*∂*u*_1_ ≠ 0 when evaluated at 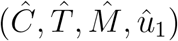, then for the curve around 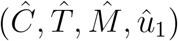, we can write *u*_1_ = *f* (*C, T, M*), where *f* is a real function.

If *T* = 0, then ∂*F/*∂*u*_1_ = *ω*_1_ *>* 0. Assuming *T* is nonzero, then ∂*F/*∂*u*_1_ = 0 only when

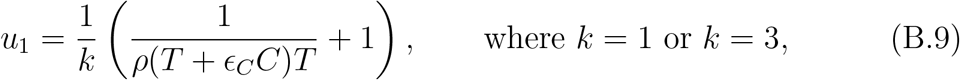

giving a condition for the uniqueness of the characterization of *u*_1_.

Minimizing the Hamiltonian (B.1) with respect to *u*_2_,

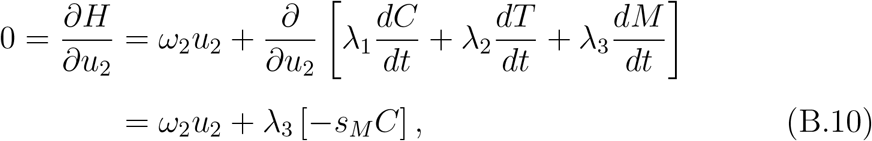

which implies that *u*_2_ is uniquely characterized by

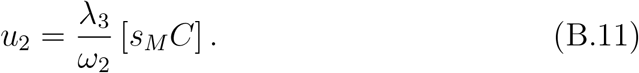

Therefore, the optimal pair 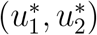 exists, and further, it is unique under (B.9).

## Appendix C. Structural Identifiability

Structural identifiability analysis investigates the ability to determine model parameters based on the structure of the model, without regard for the available data set (Guillaume et al. (2019)). If a model is structurally non-identifiable, this suggests that the model structure should be altered to be more suited for the available data, or the data type should be reconsidered to be more applicable to the model.

### Appendix C.1. Overview of the differential algebra approach

Using concepts from differential algebra (Dummit and Foote (2004)), Audoly et al. (2001) developed a method to identify the global structural identifiability of nonlinear models. This method generates input-output equation(s), which can be used to uniquely estimate parameters, and has been used to study the identifiability of several biological models (Eisenberg and Jain (2017); Eisenberg et al. (2013); Saccomani (2010); Remien et al. (2021)).

#### Definition 1.

*A parameter p*_*i*_ *is* globally *(or* uniquely) structurally identifiable *if only a single value for p*_*i*_ *results in the observed output; in other words, the model output is injective almost everywhere with respect to p*_*i*_. *In the weaker case, a parameter p*_*i*_ *is* locally *(or* non-uniquely) structurally identifiable *when a finite number of values for p*_*i*_ *generate the observed output. If a parameter is neither globally nor locally structurally identifiable, then it is* structurally non-identifiable.

#### Definition 2.

*A model is* structurally non-identifiable *if it has at least one non-identifiable parameter*, locally structurally identifiable *if all parameters are at least locally identifiable, and* globally structurally identifiable *if all parameters are globally identifiable*.

The differential algebra approach proceeds as follows. Consider the system

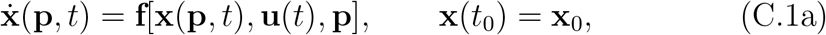

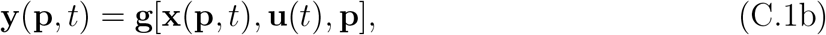

where **x, u, y**, and **p** are vectors of states, inputs, outputs, and parameters, respectively, and **f** and **g** are vectors of polynomial or rational functions in **p** and time *t*. Both the inputs, **u**, and outputs, **y**, come from experimental data. The input, **u**, represents experimental changes to the system (if any), while the output, **y**, is a measured variable, such as cell population counts.

As in Ollivier (1990), the differential ring is *R*(**p**)[**x, y, u**], where the variables are the states, input, and outputs, and where *R*(**p**) is the ring generated by the parameters. The set of differential polynomials, 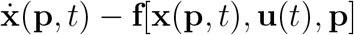 and **y**(**p**, *t*) − **g**[**x**(**p**, *t*), **u**(*t*), **p**], generate a differential ideal within *R*(**p**)[**x, y, u**].

Next, the variables are ranked to reduce computational complexity. This is, in a sense, an ordering of the variables and their derivatives within the differential ideal, just as one alphabetizes words depending on the order of the alphabet. Using this ranking, differential polynomials in the ideal can be strictly ordered to form chains, where any chain of lowest rank is called a *characteristic set*. For this method, it has been shown that the characteristic set is unique (Audoly et al. (2001); Ljung and Glad (1994)). The ranking of variables results in the characteristic set to be in triangular form. For example, if we determine that the ranking of variables is **u** *<* **y** *<* **x**, the characteristic set is

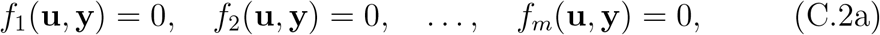

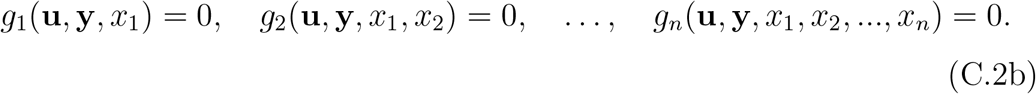

The *input-output equation(s)* are the polynomial(s) from the characteristic set which only contain the input and output variables, i.e., the polynomials *f*_*i*_. These polynomials can yield important information regarding structural identifiability because they only use variables **u** and **y**, which are explicitly known from the data.

Assuming that the variables of the input-output equation(s) do not vanish, the coefficients of *f*_*i*_ form a system of nonlinear equations expressed as multivariable polynomials with respect to the parameters, **p**. This set of coefficients is referred to as the *exhaustive summary* of the model. Parameters can be uniquely determined using these coefficients, as it is assumed that the exhaustive summary contains all structurally identifiable combinations for the model.

### Appendix C.2. Structural identifiability of the treatment-free model

Using the differential algebra approach, we investigate the structural identifiability of the model (1) without treatment.

#### Theorem 2

(Structural Identifiability). *The treatment-free model* (1), *with u*_1_ = 0 *and u*_2_ = 0, *is globally structurally identifiable from data quantifying the total number of cancer cells*.

*Proof*. The treatment-free model (1), with *u*_1_ = 0 and *u*_2_ = 0, is

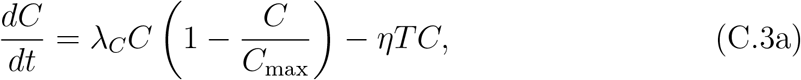

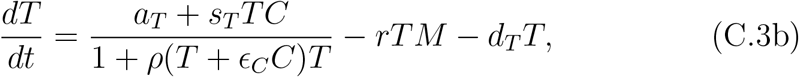

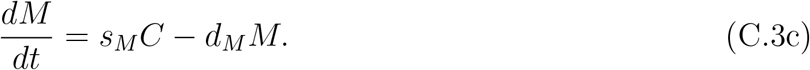

The ranking *C < M < T* is given to the variables to ensure that the input-output equation would consist of only one cell type, namely the cancer cells, *C*. The characteristic set is *f* (*C*), *g*_1_(*C, M*), and *g*_2_(*C, M, T*), and the input-output equation, *f* (*C*), contains information on model identifiability.

To determine *f* (*C*), we first solve (C.3a) for *T* and input this into (C.3b). Then, solving (C.3b) for *M*, we substitute this into (C.3c). This produces an equation which is entirely in terms of the variable, *C*, its derivatives, and the 11 positive and real parameters. We multiply this by the lowest common denominator to obtain the input-output equation, *f* (*C*), which contains 107 monomial terms.

Solving from the coefficients of the terms 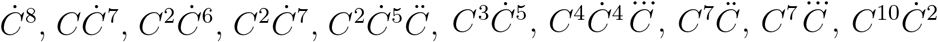, and *C*^15^, we obtain unique solutions for each of the 11 parameters. Thus, since all the individual parameters are identifiable, the treatment-free model (C.3) is globally structurally identifiable.

If we were to consider structural identifiability from data quantifying the total number of cells for all three cell types, the output would be **y** = (*C, T, M*). Thus, the input-output equations would be the entire characteristic set, which is obtained by rearranging system (C.3) in polynomial form. The corresponding calculations with data **y** = (*C, T, M*) are similar to those in proof of Theorem 2, and the model is also globally structurally identifiable in this case.

## Appendix D. Practical Identifiability

Practical identifiability analysis determines the ability to uniquely identify parameter values given a specific data set. Although a model cannot be practically identifiable unless it is structurally identifiable, structural identifiability does not imply practical identifiability (Guillaume et al. (2019)), as structural identifiability assumes that the data is noise-free and perfectly measured, which is unrealistic in practice. Even if a model is globally structurally identifiable, it can be impossible to accurately estimate parameters if the model is especially sensitive to data sparsity or measurement errors, since these data issues can mask principal features of the model dynamics (Eisenberg and Jain (2017)).

We use a sparse and noisy data set consisting of total cell counts of cancer cells, T cells, and MDSCs in murine gliomas reported in Anderson et al. (2023) to conduct practical identifiability analysis. Mice were orthotopically implanted with 35,000 KR158 high-grade glioma cells and then euthanized on days 7, 13, 20, 24, 27, and 34 following glioma implantation. Fluorescent imaging and quantification produced tumor, T cell, and MDSC cell count data for each tumor resection.

To fit the treatment-free model (C.3) to the experimental data set, we use the gradient descent method and minimize the averaged least squares error

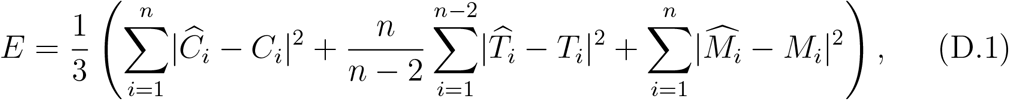

where the individual terms represent the least squares error with respect to cancer cells, T cells, and MDSCs, respectively. 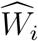 and *W*_*i*_ are the experimental and simulated data points, respectively, at the *i*^th^ time point, for *W* = *C, T, M*. The experimental data points are taken to be the averaged cell counts over multiple mice. The T cell error term is weighted by *n/*(*n* − 2) because there are two fewer data points for the T cells than the cancer cells and MDSCs.

### Appendix D.1. Fisher information matrix

Defining the estimated parameters for one data set to be the set 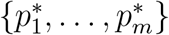, the next step is to calculate the Fisher information matrix (FIM), which is used to numerically determine local identifiability. The simplified FIM, *F*, is the Gramian of the sensitivity matrix, *X*, i.e.,

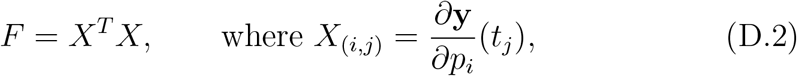

for output **y**, time points *t*_1_, …, *t*_*n*_, and parameters *p*_1_, …, *p*_*m*_. For each column *i* of *X*, we approximate the derivative of the output at each data point with respect to parameter, *p*_*i*_, at the estimate, 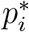. The ranks of the FIMs, *F*_*C*_, *F*_*T*_, and *F*_*M*_, calculated individually for each variable (*C, T*, and *M*) were 4, 5, and 5, respectively, suggesting that at most 5 parameters can be identified with the cell population data.

#### Appendix D.2. Sensitivity analysis

Since there are 11 parameters in the treatment-free model (C.3) but at most 5 appear to be practically identifiable, we additionally use sensitivity analysis (SA), which evaluates the influence that shifting a parameter value has on the model output, to narrow down the search for identifiable parameters. This analysis has a forward approach, in contrast to the backward approach of identifiability analysis, which determines the effect that varying data has on parameter estimates. Essentially, if a model is insensitive to a parameter, the parameter will be non-identifiable (Guillaume et al. (2019)).

In Anderson et al. (2023), sensitivity analysis of the treatment-free model (C.3) indicated that the six most sensitive parameters are *λ*_*C*_, *C*_max_, *η, ρ, r*, and *d*_*M*_. We fixed the remaining parameters to the “Best Fit” values listed in Table 1, and then re-estimated the six sensitive parameters by minimizing (D.1) using gradient descent. The estimated parameter values are listed in Table D.2.

**Table D.2:** Re-estimation of the six most sensitive parameter values of the treatment-free model (C.3) using murine glioma data from Anderson et al. (2023). The remaining model parameters were fixed to the “Best Fit” values listed in Table 1, and then estimation was performed by minimizing (D.1) using the gradient descent method.

| Parameter | Value | Units |
| --- | --- | --- |
| $\lambda_C$ | 0.483 | day<sup>-1</sup> |
| $C_{\max}$ | $1.17 \times 10^8$ | cell |
| $\eta$ | $1.62 \times 10^{-7}$ | day<sup>-1</sup> cell<sup>-1</sup> |
| $\rho$ | 2.15 | cell<sup>-2</sup> |
| $r$ | $1.29 \times 10^{-7}$ | day<sup>-1</sup> cell<sup>-1</sup> |
| $d_M$ | 0.217 | day<sup>-1</sup> |

#### Appendix D.3. Profile likelihood

We generate profile likelihoods for the six parameters *λ*_*C*_, *C*_max_, *η, ρ, r*, and *d*_*M*_ by first selecting one parameter, *p*_*i*_, and generating 20 uniformly distributed random samples of this parameter within the neighborhood of its parameter estimate, 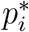. Then, using the gradient descent method and minimizing (D.1), we fit the remaining parameters to the data. Minimizing the least squares cost function (D.1) is equivalent to maximizing the likelihood function (Renardy et al. (2022); Eisenberg and Jain (2017)). The error (D.1) for each parameter *p*_*i*_ is plotted in Figure D.7, and its estimated parameter value from Table D.2, 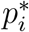, is marked as a red point.

Similar to Raue et al. (2009) and Eisenberg and Jain (2017), we calculate confidence intervals at a level of significance *α* = 0.05 for each of the tested parameters *p*_*i*_ by calculating the threshold

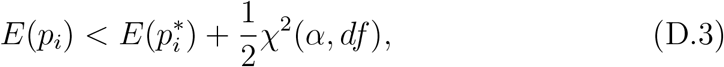

where *χ* is the chi-squared distribution for *df* = 6 parameters. The 95% confidence intervals are indicated in Figure D.7 with red dashed lines. Narrower confidence intervals suggest higher confidence in the parameter estimate, 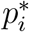.

**Figure D.7:**
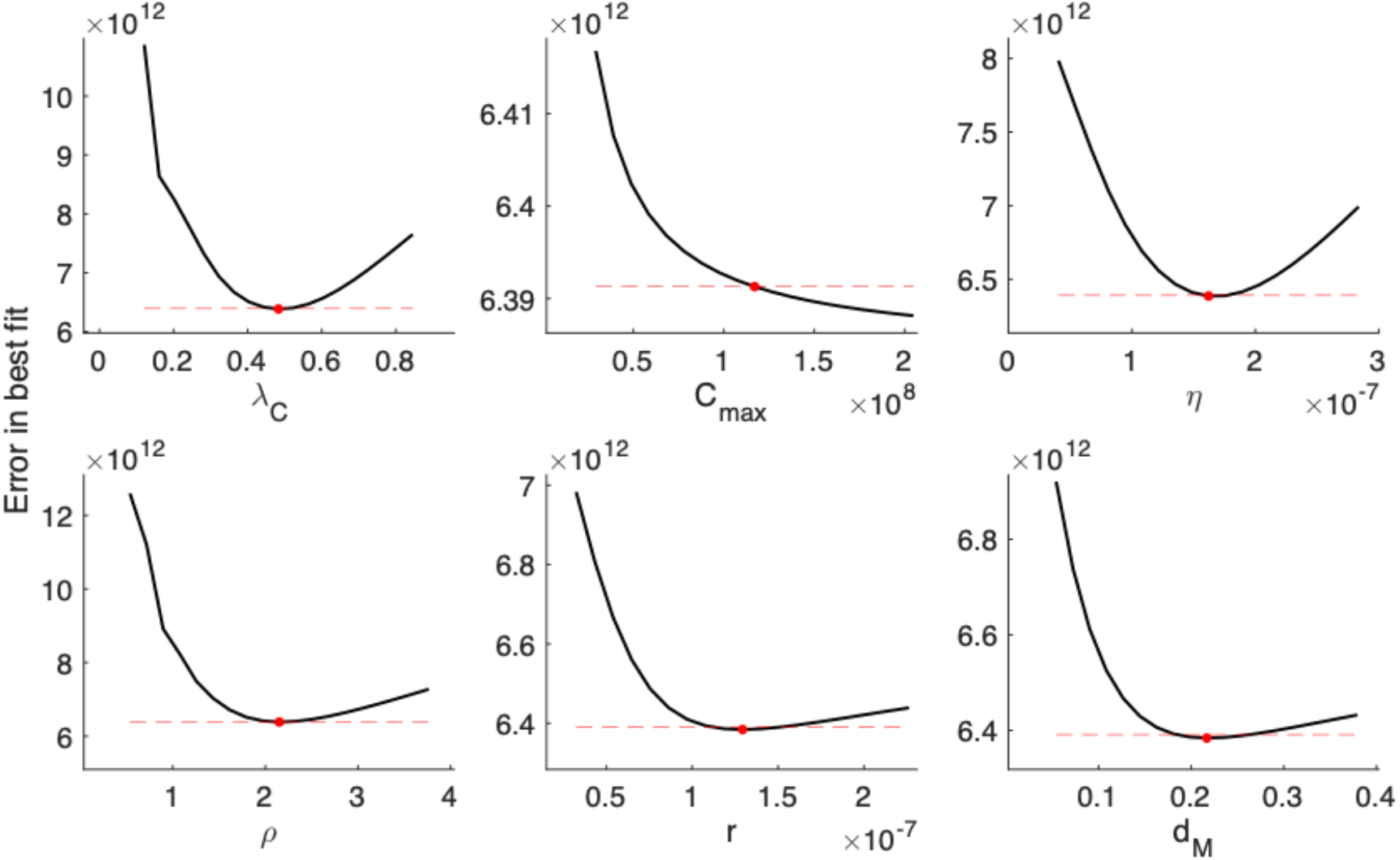
Profile likelihoods for the tumor growth rate (*λ*_*C*_), the tumor carrying capacity (*C*_max_), the tumor kill rate by T cells (*η*), the T cell inhibition rates by PD-L1-PD-1 (*ρ*) and by MDSCs (*r*), and the MDSC death rate (*d*_*M*_). Curves were determined by fixing parameter *p*_*i*_ at points along the domain and then applying the gradient descent method with murine data from Anderson et al. (2023) to find the minimal least squares error (D.1). The red point identifies the overall best fitting parameter set *{λ*_*C*_, *C*_max_, *η, ρ, r, d*_*M*_ *}* (given in Table D.2), and the red dashed line indicates the 95% confidence interval. The lack of a relative minimum suggests that *C*_max_ is not practically identifiable, while *λ*_*C*_, *η, ρ, r*, and *d*_*M*_ are practically identifiable.

Profile likelihoods of identifiable parameters exhibit distinct local minima, while structurally non-identifiable parameters have flat profile likelihoods and practically non-identifiable parameters have shallow minima. Thus, a profile likelihood which does not display a distinct minimum suggests that the parameter is non-identifiable because many values in the parameter domain generate a similar error. Figure D.7 indicates that the cell population data can practically identify the following parameters: the tumor growth rate (*λ*_*C*_), the tumor kill rate by T cells (*η*), the T cell inhibition rates by PDL1-PD-1 (*ρ*) and by MDSCs (*r*), and the MDSC death rate (*d*_*M*_). This is consistent with the FIM results (Appendix D.1), which implied that there were five practically identifiable parameters.

We conclude that *λ*_*C*_, *η, ρ, r*, and *d*_*M*_ are suitable targets for treatment personalization, while the remaining parameters are practically nonidentifiable. Numerical simulations using parameter estimates from Table D.2 are plotted in Figure D.8 alongside the experimental data from Anderson et al. (2023).

**Figure D.8:**
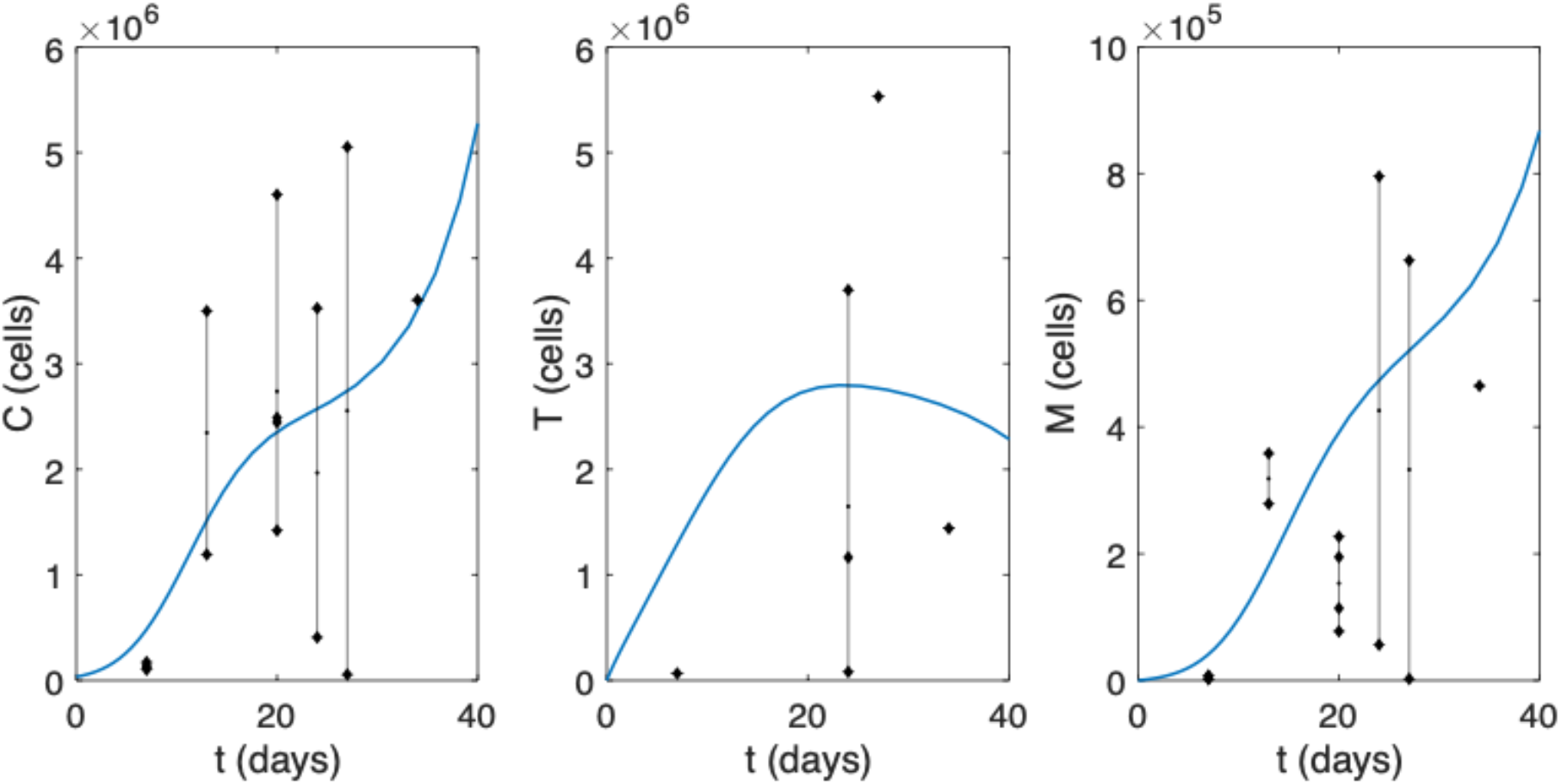
Numerical simulations of tumor cells (*C*), T cells (*T*), and MDSCs (*M*) using the parameter estimates from Table D.2. Simulations are plotted alongside murine data described in Anderson et al. (2023).

## Appendix E. Parameter summary statistics for Section 6.1

**Table E.3:** Parameter summary statistics for non-treated bladder and breast tumors in mice from Tu et al. (2020). Parameters were obtained by accepting 20,000 parameter sets of the lowest error after testing 100,000 sets using the ABC rejection method.

| Parameter |  | Bladder tumor | Breast tumor |
| --- | --- | --- | --- |
| $\lambda_C$ | Best Fit | 0.305 | 0.472 |
|  | Mean | 0.162 | 0.304 |
|  | Median | 0.148 | 0.323 |
|  | Mode | 0.159 | 0.406 |
|  | SD | 0.104 | 0.129 |
| $C_{\max}$ | Best Fit | $6.36 \times 10^7$ | $5.81 \times 10^7$ |
| | Mean | $2.43 \times 10^8$ | $2.44 \times 10^8$ |
| | Median | $2.39 \times 10^8$ | $2.39 \times 10^8$ |
| | Mode | $4.68 \times 10^7$ | $4.70 \times 10^7$ |
| | SD | $1.45 \times 10^8$ | $1.44 \times 10^8$ |
| $\eta$ | Best Fit | $5.46 \times 10^{-7}$ | $1.19 \times 10^{-8}$ |
| | Mean | $3.81 \times 10^{-7}$ | $1.65 \times 10^{-7}$ |
| | Median | $3.29 \times 10^{-7}$ | $1.28 \times 10^{-7}$ |
| | Mode | $9.67 \times 10^{-8}$ | $2.37 \times 10^{-8}$ |
| | SD | $2.71 \times 10^{-7}$ | $1.39 \times 10^{-7}$ |
| $\rho$ | Best Fit | 0.451 | 0.478 |
|  | Mean | 0.310 | 0.307 |
|  | Median | 0.324 | 0.323 |
|  | Mode | 0.450 | 0.432 |
|  | SD | 0.124 | 0.129 |
| $r$ | Best Fit | $2.58 \times 10^{-5}$ | $3.13 \times 10^{-5}$ |
| | Mean | $5.90 \times 10^{-5}$ | $5.73 \times 10^{-5}$ |
| | Median | $6.18 \times 10^{-5}$ | $6.01 \times 10^{-5}$ |
| | Mode | $8.94 \times 10^{-5}$ | $8.46 \times 10^{-5}$ |
| | SD | $2.63 \times 10^{-5}$ | $2.74 \times 10^{-5}$ |
| $d_M$ | Best Fit | 0.332 | 0.336 |
|  | Mean | 0.227 | 0.226 |
|  | Median | 0.215 | 0.215 |
|  | Mode | 0.0664 | 0.0603 |
|  | SD | 0.145 | 0.144 |

**Table E.4:** Parameter summary statistics regarding treatment efficacy and decay. Parameters were obtained by accepting 20,000 parameter sets of the lowest error after testing 100,000 sets using the ABC rejection method with bladder and breast tumor data from Tu et al. (2020). The murine tumors were treated with either anti-PD-1 and/or a CCR2 antagonist.

| **Bladder tumor** |  |  |  |  |
| --- | --- | --- | --- | --- |
| Parameter |  | Anti-PD-1<br>monotherapy | CCR2 antagonist<br>monotherapy | Anti-PD-1 +<br>CCR2 antagonist |
| $\gamma_1$ | Best Fit | $6.21 \times 10^3$ | – | $1.61 \times 10^4$ |
| | Mean | $1.42 \times 10^4$ | – | $1.16 \times 10^4$ |
| | Median | $1.52 \times 10^4$ | – | $1.26 \times 10^4$ |
| | Mode | $1.80 \times 10^4$ | – | $1.73 \times 10^4$ |
| | SD | $3.96 \times 10^3$ | – | $5.29 \times 10^3$ |
| $d_1$ | Best Fit | 1.01 | – | 1.69 |
|  | Mean | 2.94 | – | 4.17 |
|  | Median | 2.58 | – | 3.29 |
|  | Mode | 1.69 | – | 1.73 |
|  | SD | 1.47 | – | 2.60 |
| $\gamma_2$ | Best Fit | – | $2.00 \times 10^4$ | $1.80 \times 10^4$ |
| | Mean | – | $1.41 \times 10^4$ | $1.31 \times 10^4$ |
| | Median | – | $1.48 \times 10^4$ | $1.40 \times 10^4$ |
| | Mode | – | $1.84 \times 10^4$ | $1.81 \times 10^4$ |
| | SD | – | $4.07 \times 10^3$ | $4.97 \times 10^3$ |
| $d_2$ | Best Fit | – | 2.24 | 2.09 |
|  | Mean | – | 2.05 | 3.78 |
|  | Median | – | 2.90 | 2.90 |
|  | Mode | – | 2.19 | 1.84 |
|  | SD | – | 0.982 | 2.25 |
| **Breast tumor** |  |  |  |  |
| Parameter |  | Anti-PD-1<br>monotherapy | CCR2 antagonist<br>monotherapy | Anti-PD-1 +<br>CCR2 antagonist |
| $\gamma_1$ | Best Fit | $1.89 \times 10^4$ | – | $1.49 \times 10^4$ |
| | Mean | $1.43 \times 10^4$ | – | $1.20 \times 10^4$ |
| | Median | $1.53 \times 10^4$ | – | $1.30 \times 10^4$ |
| | Mode | $1.80 \times 10^4$ | – | $1.74 \times 10^4$ |
| | SD | $3.80 \times 10^3$ | – | $5.12 \times 10^3$ |
| $d_1$ | Best Fit | 1.00 | – | 1.20 |
|  | Mean | 3.06 | – | 4.10 |
|  | Median | 2.69 | – | 3.26 |
|  | Mode | 1.80 | – | 1.76 |
|  | SD | 1.55 | – | 2.55 |
| $\gamma_2$ | Best Fit | – | $1.99 \times 10^4$ | $1.77 \times 10^4$ |
| | Mean | – | $1.45 \times 10^4$ | $1.31 \times 10^4$ |
| | Median | – | $1.51 \times 10^4$ | $1.41 \times 10^4$ |
| | Mode | – | $1.85 \times 10^4$ | $1.81 \times 10^4$ |
| | SD | – | $3.83 \times 10^3$ | $4.99 \times 10^3$ |
| $d_2$ | Best Fit | – | 2.13 | 1.54 |
|  | Mean | – | 3.14 | 3.88 |
|  | Median | – | 2.98 | 2.97 |
|  | Mode | – | 2.18 | 2.00 |
|  | SD | – | 1.03 | 2.30 |

